# Identifying fungal leaf spots on oilseed rape: could pictures help?

**DOI:** 10.1101/2022.10.21.513129

**Authors:** L. Bousset, M. Ermel, B. Bammé, A. Penaud, J. Carpezat, MH. Balesdent, V. Laval, M. Palerme, N. Parisey

## Abstract

From sowing in late summer until harvest in following summer, oilseed rape can be infected by several fungi, which foliar symptoms (leaf spots) coexist on the crop. Training an expert at their identification is quick for the typical symptoms with characteristic appearance. However, in many cases the size, colour and morphology are similar and for the atypical symptoms, there is a risk of confusion or in-decidability. Also, scouting the fields for expert training is not possible at all seasons and all diseases might not be seen in all years and all places. The aim of our study was to produce large sets of pictures annotated by several experts, from which tables illustrating the diversity of symptom appearance were chosen. These tables will enable assistance to diagnostic and expert training.

## Introduction

A critical challenge in plant pathology and epidemiology is to design and implement durable crop protection strategies against pathogens. On annual crops, many plant diseases have cyclic epidemics and their dynamics are highly influenced by both temporal and spatial discontinuities, either induced by the climate (e.g. seasonality) or by human actions (e.g. sowing and harvesting) (Zadoks & Schein 1979; Bousset & Chèvre 2013). Therefore, on the one hand each disease might not be present during the whole cropping season of the host. On the other hand, a given host can simultaneously face several pathogens. Precise identification of each causal agent is needed to deploy proper crop protection strategies.

In Western Europe, oilseed rape is grown as a winter crop, sown in late summer and harvested in the following summer. During this extended period of time, several foliar pests and diseases can infect leaves and cause co-occurring symptoms on the crop. Identification of the causal agents is at stake for plant pathologists, breeders and extension service agents. Training an expert at their identification is quick and easy for typical symptoms with characteristic appearance. Training can be done in the field with real symptoms at seasons when symptom occur, or using symptom images. However, in many cases the size, colour and morphology are similar between different species, because for each of the species the symptom aspect is altered by several factors. The plant genotype, potentially with complete or partial resistance, can render the interaction less compatible, with plant defence reactions inducing e.g. darkening around lesions. Also, some of the symptom characteristics, such as darkening or yellow halo, can be convergent between two or more pathogens and bring confusion to the diagnostic. Finally, the physiological state of the plant tissues can also have an impact on the symptoms. So in many cases the size, colour and morphology of the symptoms can be similar although the plant is infected by different fungal species and there is a risk of confusion or indecidability between several symptom-causing pathogens.

In this study we aimed at sampling the diversity of symptom appearance for foliar diseases on oilseed rape during the cropping season, with multiexpert annotation. We gathered a community of plant pathologists more or less trained to the diagnostic of foliar diseases of oilseed rape in order to build a database of consensual or problematic symptoms. Standardized pictures of a range of symptoms were taken, independently annotated and classified in different categories depending on experts’ answers. This enabled us to build a validated set of “typical” and “atypical” symptom pictures for six foliar fungal pathogen species. Finally, this work enabled us to list the phenotypic traits that help distinguishing each of these species. From this diversity, illustrations are provided in addition to sets of distinctive criteria as material to help diagnostic or train experts.

## Materials and methods

### Experimental fields, collection of leaf symptoms and handling of leaves

Symptomatic leaves were collected from winter oilseed rape fields in cropping seasons 2018-2019 to 2020-2021 on plants ranging from cotyledon to pod formation growth stage (Table 1). To encompass the varietal diversity of winter oilseed rape, we sampled variety testing plots, plant breeding nursery, both located on the INRAE UE La Motte experimental station in Le Rheu (48.1°N, 1.5°W), in Brittany and scouted farmers’fields within 40 km around. The climate of Britany is oceanic and most of the oilseed rape foliar fungal diseases are observed each year. To expand the range of soil and cropping conditions, samples at Grignon (48.9°N, 1.9°E) from INRAe BIOGER and TERRES INOVIA, in Ile-de-France were added. Leaves sampled were either typical i.e. with causal agent identified at first sight and atypical i.e. identification not straightforward. While scouting the fields, leaves were harvested altogether in bags. In the laboratory, leaves were arranged in buckets with petioles dipped in water so that they would keep fresh overnight. Leaves were further processed either the same or the following day by imaging the leaf lesions.

**Table 1:** Numbers of leaf segments sampled depending on the year and the growth stage.

|  | All leaf fragments |
| --- | --- |
| **Season - Month** |  |
| 2018-2019 Mar. | 306 |
| 2019-2020 Oct. | 114 |
| 2019-2020 Nov. | 46 |
| 2019-2020 Feb. | 169 |
| 2019-2020 Mar. | 239 |
| 2020-2021 Nov. | 63 |
| **Growth stage** |  |
| Cotyledon | 67 |
| Leaf rosette | 156 |
| Stem elongation | 178 |
| Flowering | 456 |
| Pod formation | 80 |
| **Total** | **937** |

### Image acquisition and pre-processing

The first step was to image the leaf lesion environment on the leaf, first taking pictures of the whole area on the recto and verso sides, then taking a third picture of the recto using a red gasket placed to circle one isolated leaf lesion. Using scissors, the second step was to cut a leaf portion containing only one leaf lesion. A pair of standardised pictures on the recto and the verso side of this fragment were taken on a blue background (PVC sheet Lastolite Colormatt electric blue), together with a label indicating sample name and a second label indicating either recto or verso, a ruler and a standardised colour test pattern with white grey and black. Two workshop led hand lights were placed on both sides of the leaf fragment within the lower 45° angle. Pictures were taken with a Nikon D5200 with an AF-S DX Micro Nikkor 40mm 1:2.8G lens, on a self-assembled stand, with a wired remote control. Aperture was set at F14 for maximal depth of field, iso 125, daylight white balance. Pictures were saved as RGB images with a resolution of 6000 x 4000 pixels. Picture pre-processing consisted in reading the barcodes to rename the files. The corresponding leaf fragments were kept frozen at −20°C.

### Multi-expert annotation on a web interface

A set of 6 pathologists familiar with oilseed rape but not with each of the fungal diseases was assembled. Each of the expert had to annotate each picture on a web interface, having access to the other members’ annotations only after having completed the duty. The set of 5 pictures of each symptom (general environment recto and verso, leaf lesion circled, leaf fragment recto and verso) was available with possibility to zoom. Each symptom was annotated as follows (Fig. 1). The first question was about confidence, with 3 levels (“certain”; “I hesitate” or “I don’t know”). When “certain”, the number of symptoms enables to set aside symptomless fragments; finally if symptomatic, tick boxes were available for 13 distinct causal agents. When “hesitating”, too complex images (more than one symptom, some of which uncertain) are set aside; distinction is made between on the one hand the identification of one species but atypical symptom appearance; on the other hand hesitation between several species. When the expert “can’t tell”, the symptom is classified unidentified. For the symptomatic leaf fragments, tick boxes enable “and” and “or” selection of 9 fungal diseases, liquid fertilisation, insect, virus and bacterial damage.

**Figure 1:**
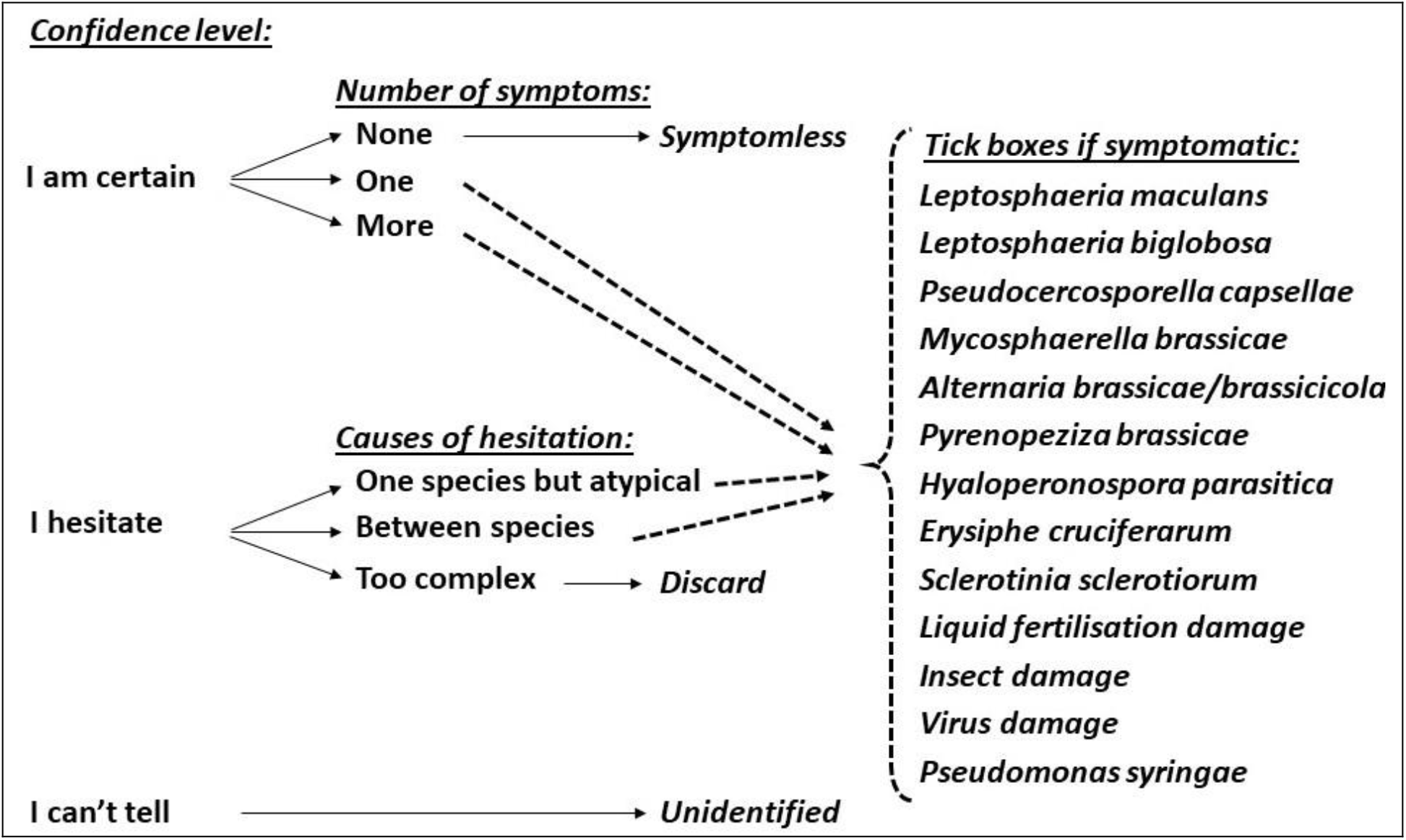
Decision key for the annotation by experts on the web interface depending on expert’s confidence level, numbers of symptoms seen, causes of hesitation and causal species.

### Set of criteria and picture selection for the figures

Following annotation of the whole dataset, the experts devised sets of criteria distinctive for each of the fungal species frequently encountered on oilseed rape in France. They selected subsets of pictures on the one hand with typical symptoms (respecting most of the distinctive criteria); on the other hand with atypical symptoms (variants not respecting all criteria).

## Results

We produced a database of 937 symptoms standardized image recto-verso pairs from a diversity of varieties and sampling times across 2019, 2020 and 2021. The annotation step distinguished between contrasted situations of expertise ranging from simple consensual cases to disagreement or lack of identification confirming the difficulty of visual and photographic diagnostic.

The 6 experts agreed on sets of distinctive criteria for each of the 6 species frequently encountered on oilseed rape in France (Fig. 2; FigS1 higher quality pictures).

**Figure 2:**
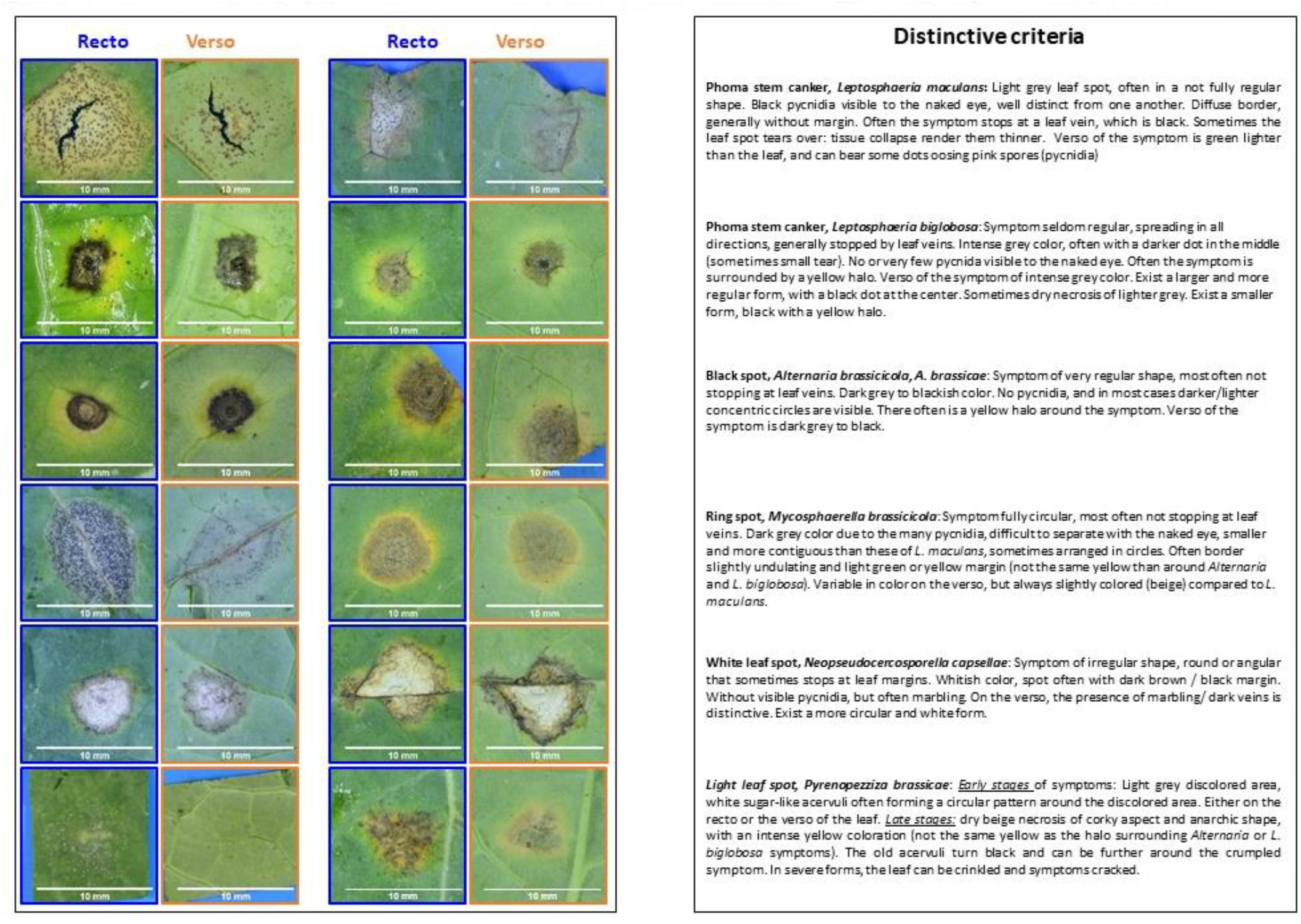
Sets of criteria distinctive for each of the six species (right panel) and corresponding illustrative recto-verso image pairs for two symptoms per species.

For each of these 6 species, the 6 experts selected subsets of pictures on the one hand with typical symptoms (respecting most of the distinctive criteria); on the other hand with atypical symptoms (variants not respecting all criteria). Figures were produces for phoma stem canker *Leptosphaeria maculans* (Fig. 3 & Fig. S2 higher quality pictures) and *Leptosphaeria biglobosa* (Fig. 4 & Fig. S3 higher quality pictures); for white leaf spot *Neopseudocercosporella capsellae* (Fig. 5 & Fig. S4 higher quality pictures); for ring spot *Mycosphaerella brassicae* (Fig. 6 & Fig. S5 higher quality pictures); for black spot *Alternaria brassicae* and *A. brassicicola* (Fig. 7 & Fig. S6 higher quality pictures) and light leaf spot *Pyrenopezziza brassicae* (Fig. 8 & Fig. S7 higher quality pictures).

**Figure 3:**
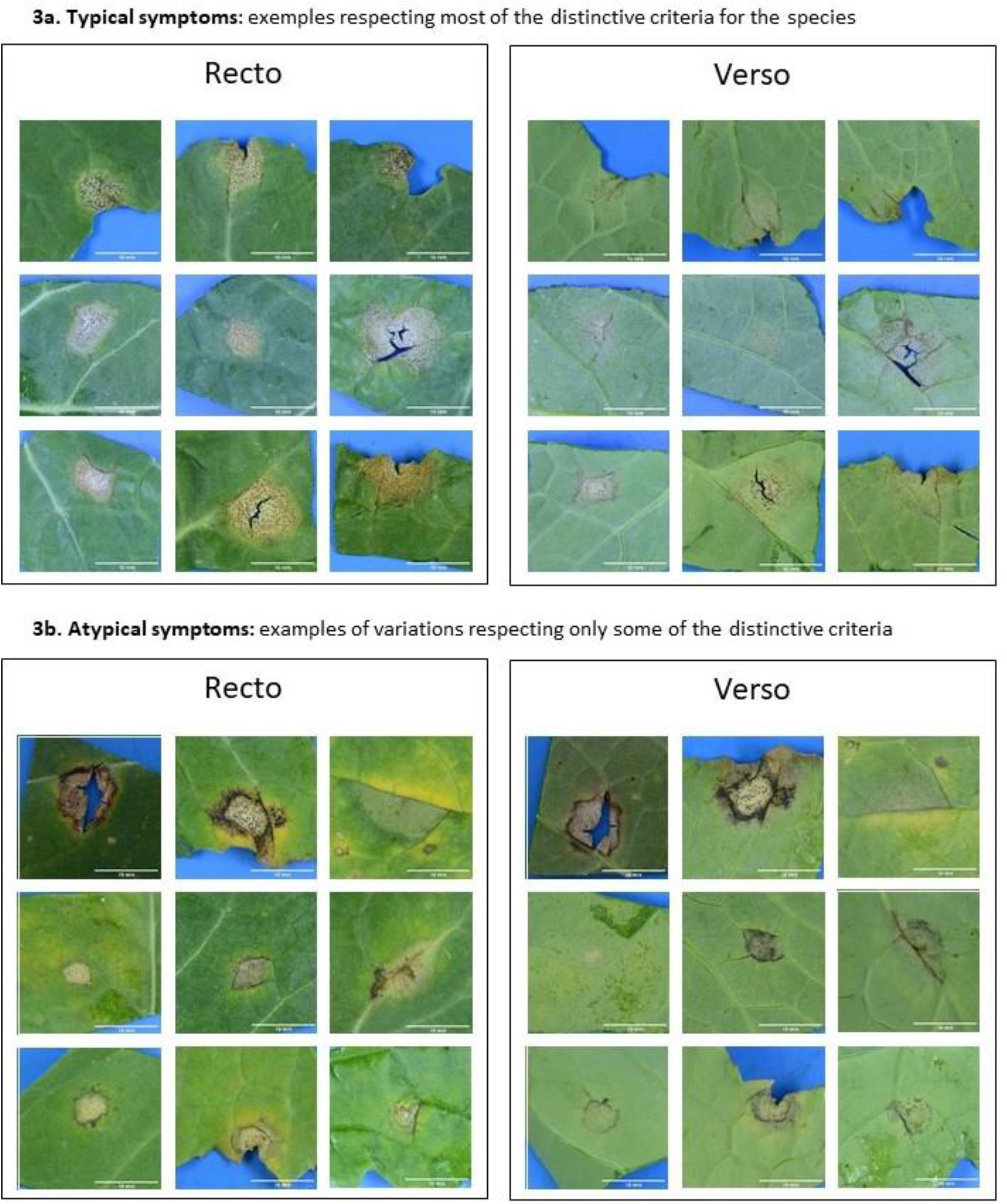
Phoma stem canker *Leptosphaeria maculans*

**Figure 4:**
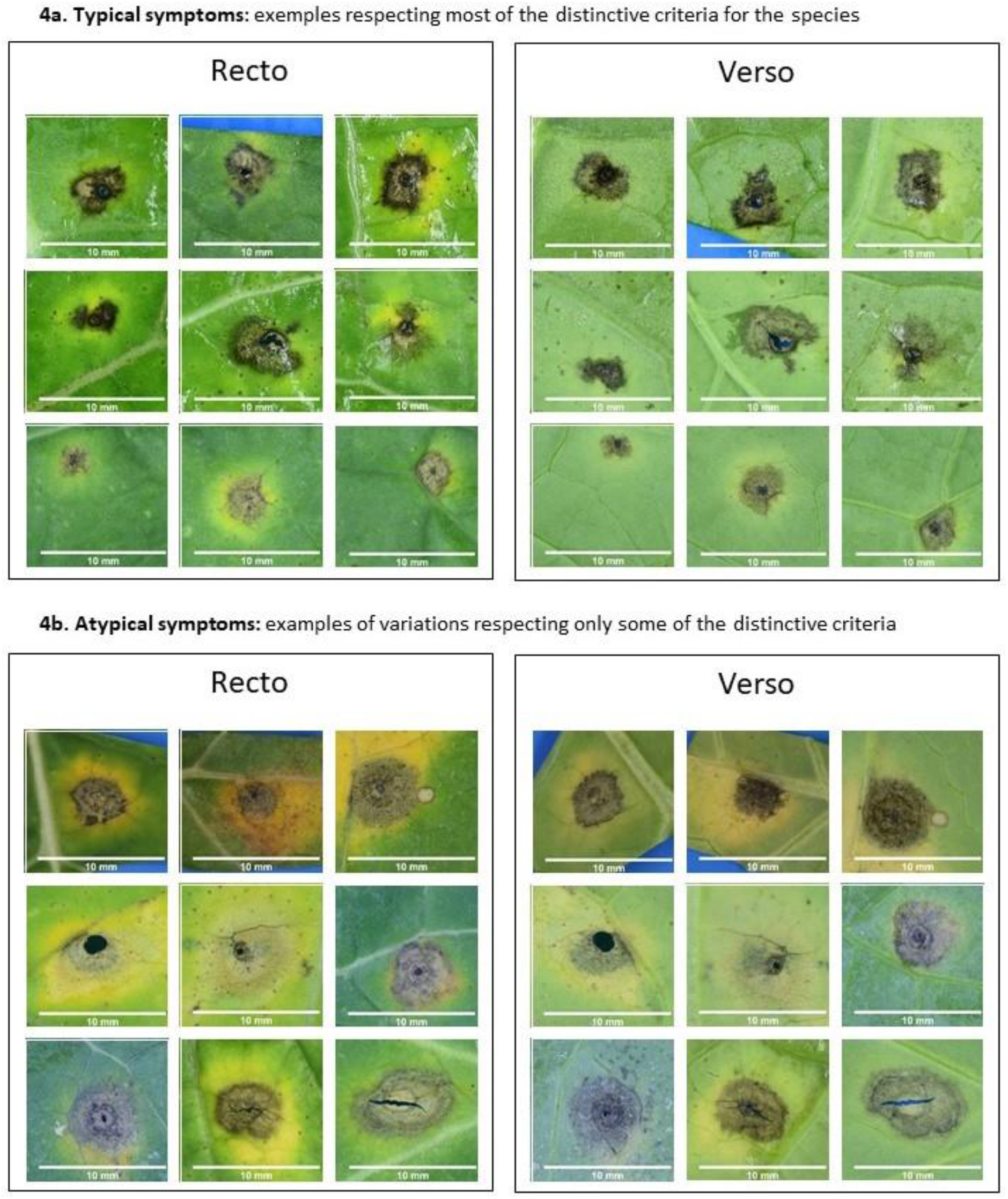
Phoma stem canker *Leptosphaeria biglobosa*

**Figure 5:**
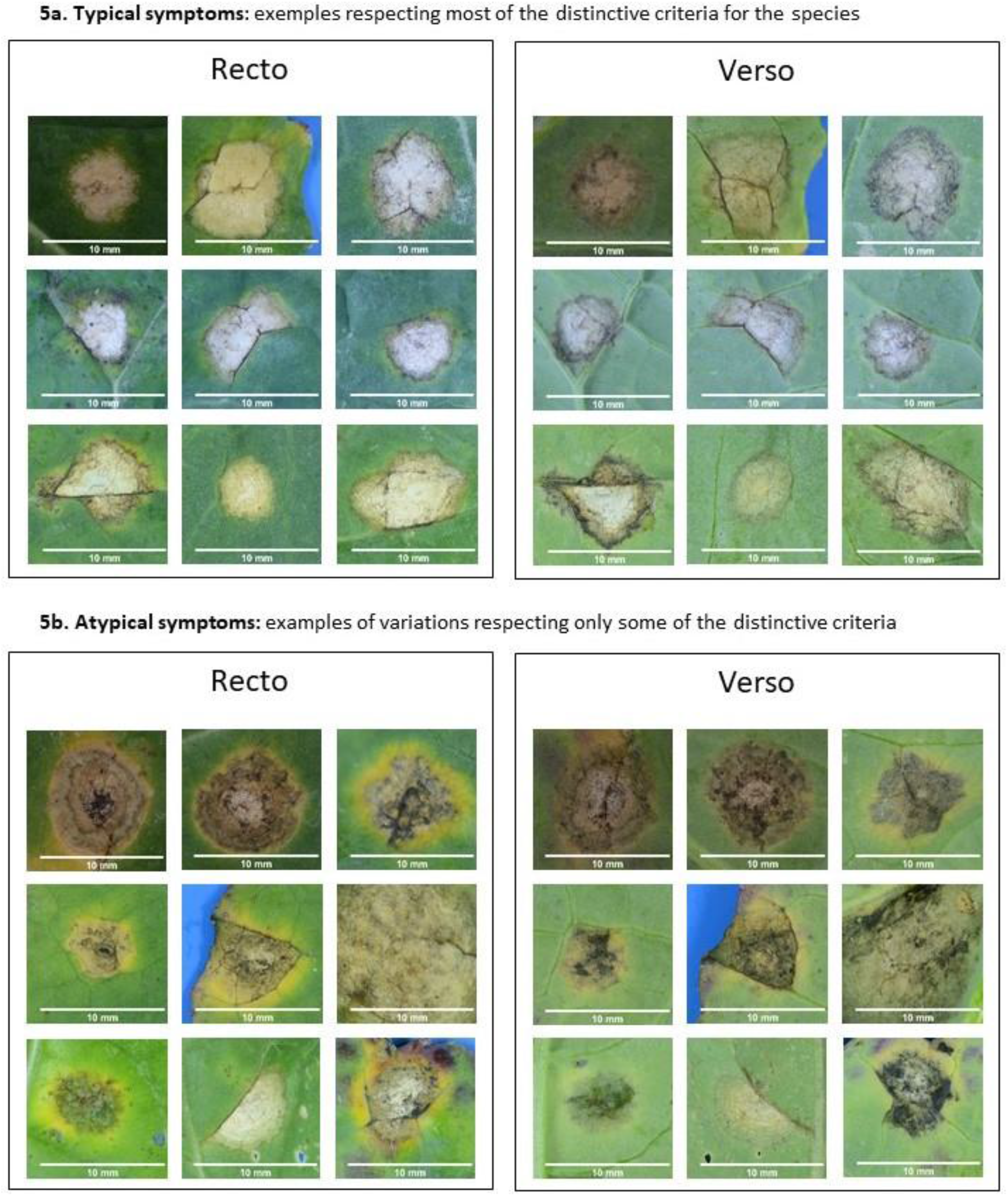
White leaf spot *Neopseudocercosporella capsellae*

**Figure 6:**
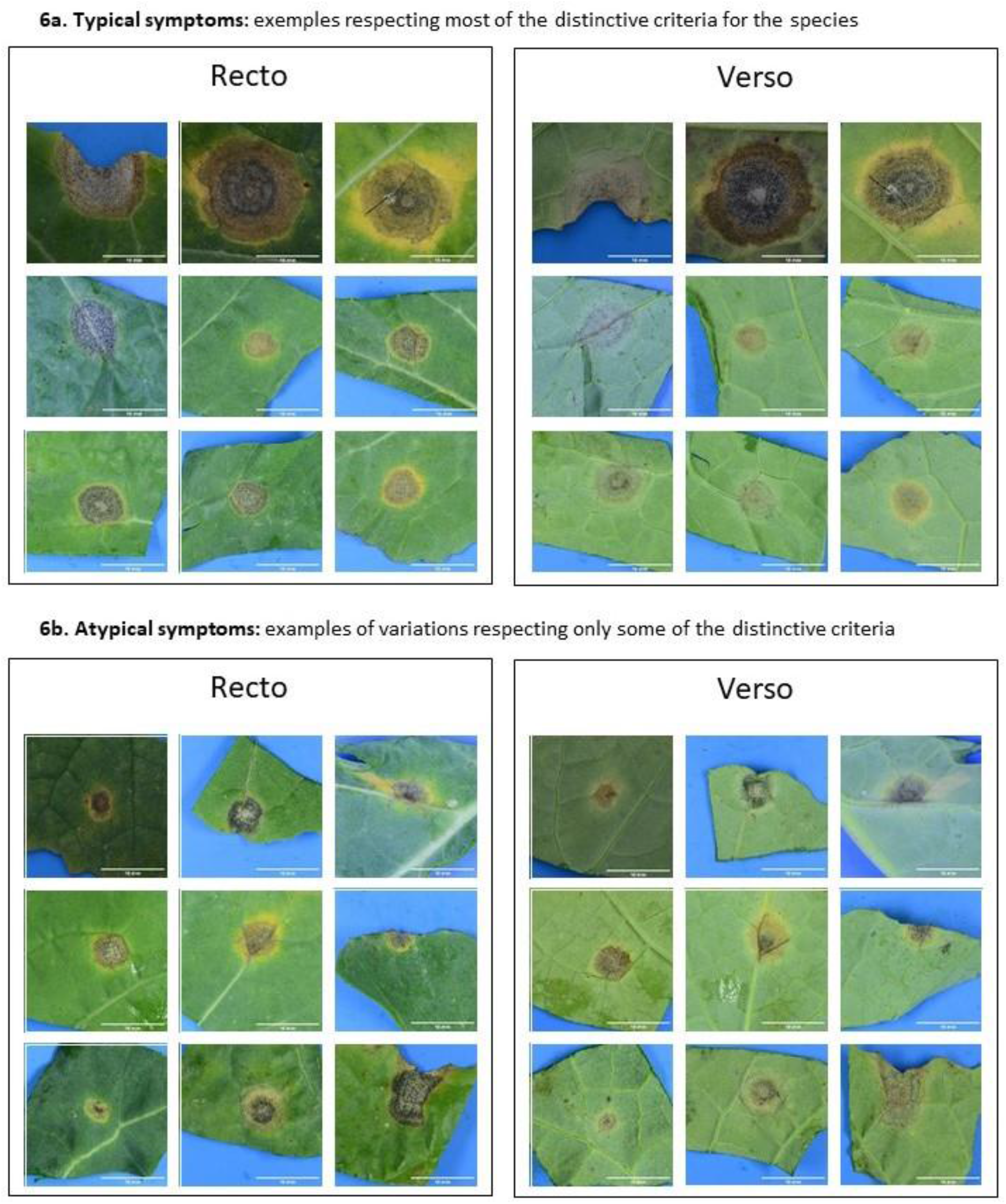
Ring spot *Mycosphaerella brassicae*

**Figure 7:**
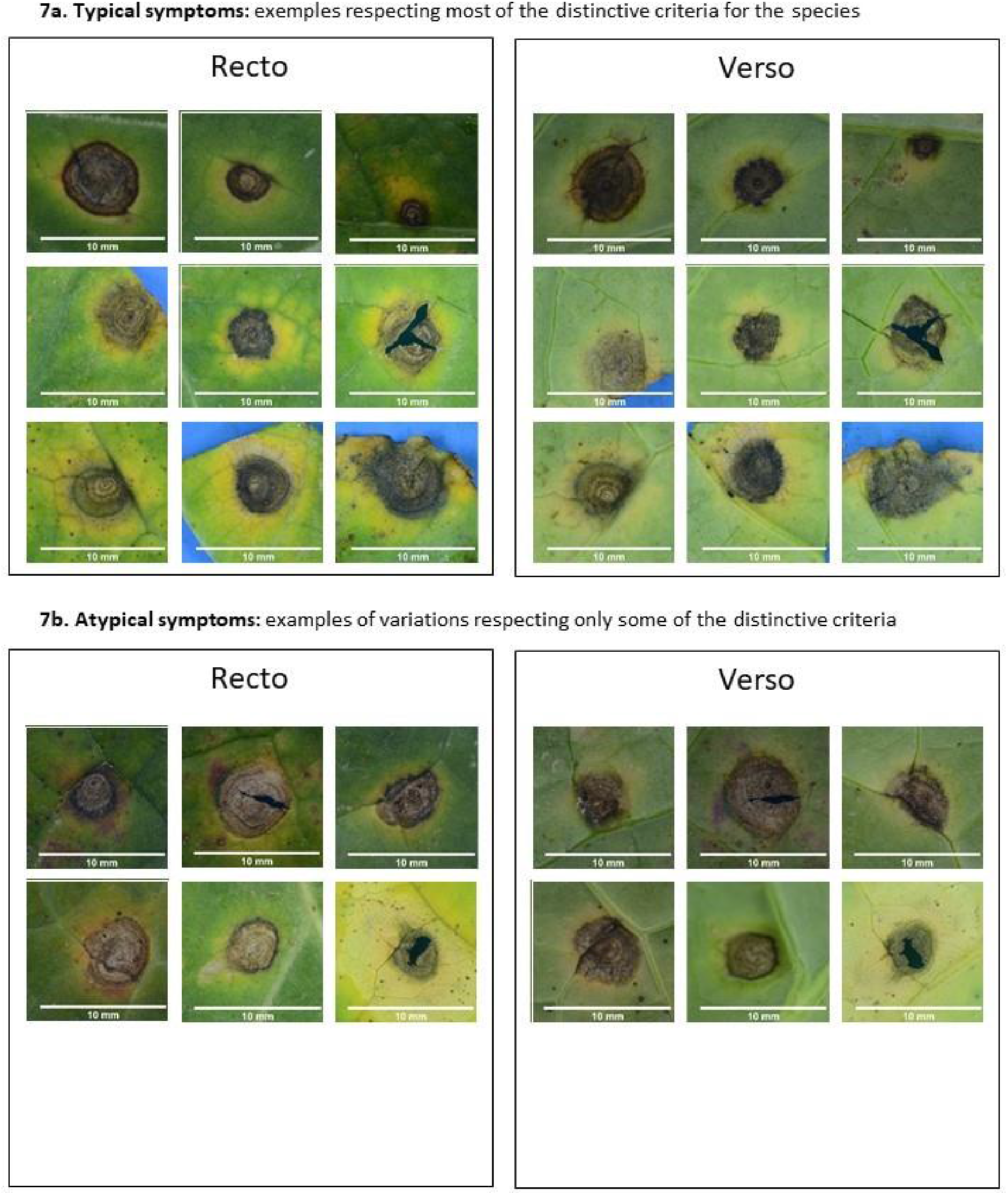
Black spot *Alternaria brassicae* and *A. brassicicola*

**Figure 8:**
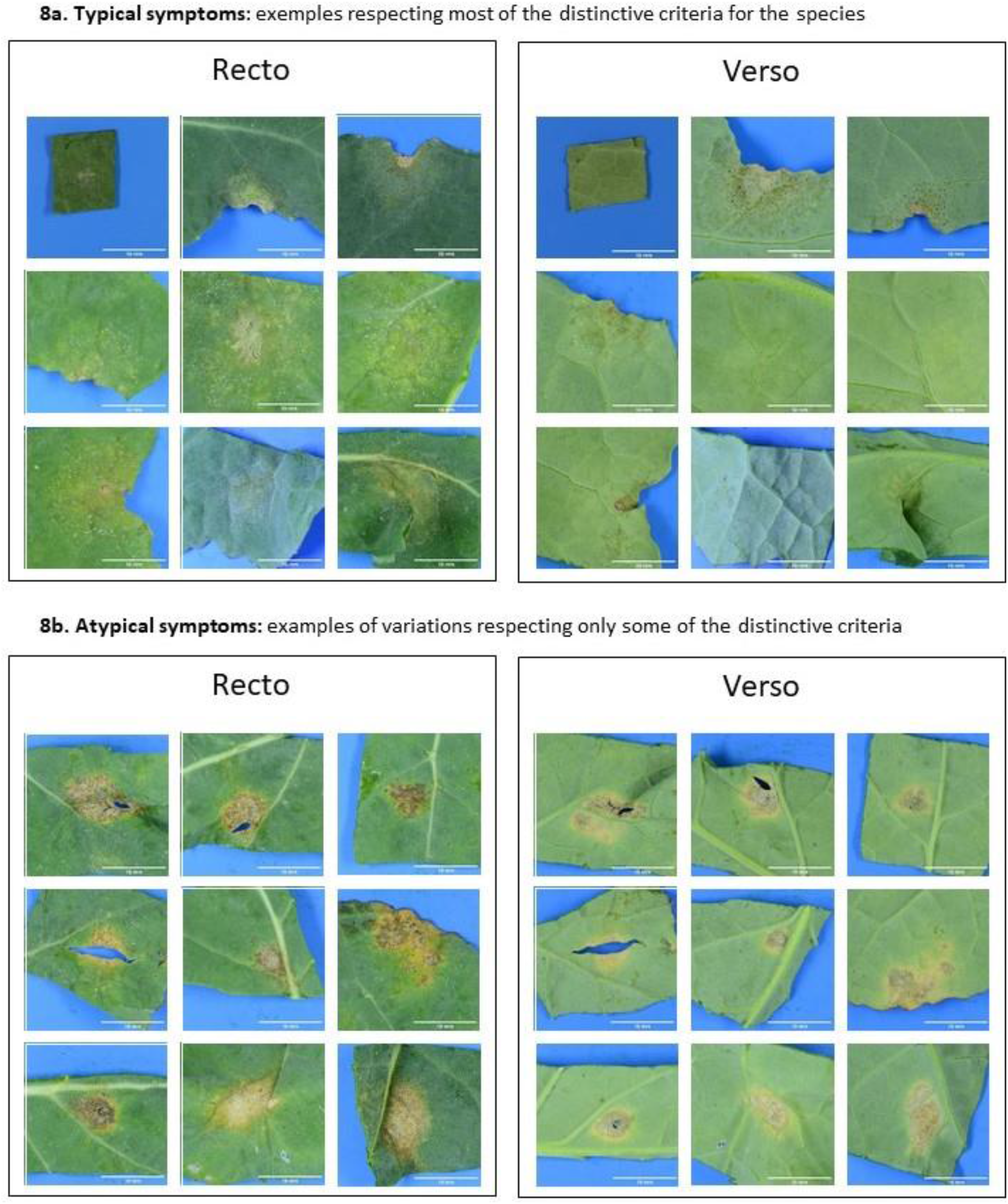
Light leaf spot *Pyrenopezziza brassicae*

## Discussion

We confirmed that along the growth of oilseed rape several fungal diseases co-occur on leaves. This might not be the case in all regions, but the oceanic climate of Brittany allowed observing all the major diseases.

The experts were not able to identify distinctive criteria that would be respected by all the symptoms of a given species and not frequently or occasionally shared with other species. However, combining several criteria for selection or exclusion allows to produce guidelines for typical symptoms. For the atypical symptoms (variations respecting only some of the distinctive criteria, the availability of standardized pictures will be of great help for the reader.

In addition to the current illustrative figures, the availability of our picture database will enable developing computer applications to train experts. The annotated images can be selectively presented to candidates, rated and performance evaluated before going to the field.

As the leaf fragments corresponding to leaf fragments in the image database were kept frozen, the prospect is open to confirm expert visual diagnostic by DNA molecular characterisation.

## Acknowledgements

We thank UE La Motte and surrounding farmers for the cultivation of oilseed rape fields as well as the INRAe experimental unit in Grignon (Christophe Montagnier). We thank Yannick Lucas for technical assistance. This work benefited from the financial support of INRAe - the French National Institute for Agronomical Research, and from CASDAR project C2018 – 11 (Atipical). The “Effectors and Pathogenesis of *L. maculans*” group benefits from the support of Saclay Plant Sciences-SPS (ANR-17-EUR-0007).

## Supplementary Materials

In Supplementary figures Fig. S1 to Fig. S7, the contents of Figure 2 to Figure 8 are provided with higher quality pictures.

