## SupplementaryMaterials for "Identifying fungal leaf spots on oilseed rape: could pictures help?"

**Figure S1:** *Sets of criteria distinctive for each of the six species (right panel) and corresponding illustrative recto-verso image pairs for two symptoms per species.*

**Figure S2:** Phoma stem canker *Leptosphaeria maculans*

**Figure S3:** Phoma stem canker *Leptosphaeria biglobosa*

**Figure S4:** White leaf spot *Neopseudocercosporella capsellae*

**Figure S5:** Ring spot *Mycosphaerella brassicae*

**Figure S6:** Black spot *Alternaria brassicae* and *A. brassicicola*

**Figure Sè: L**ight leaf spot *Pyrenopezziza brassicae*

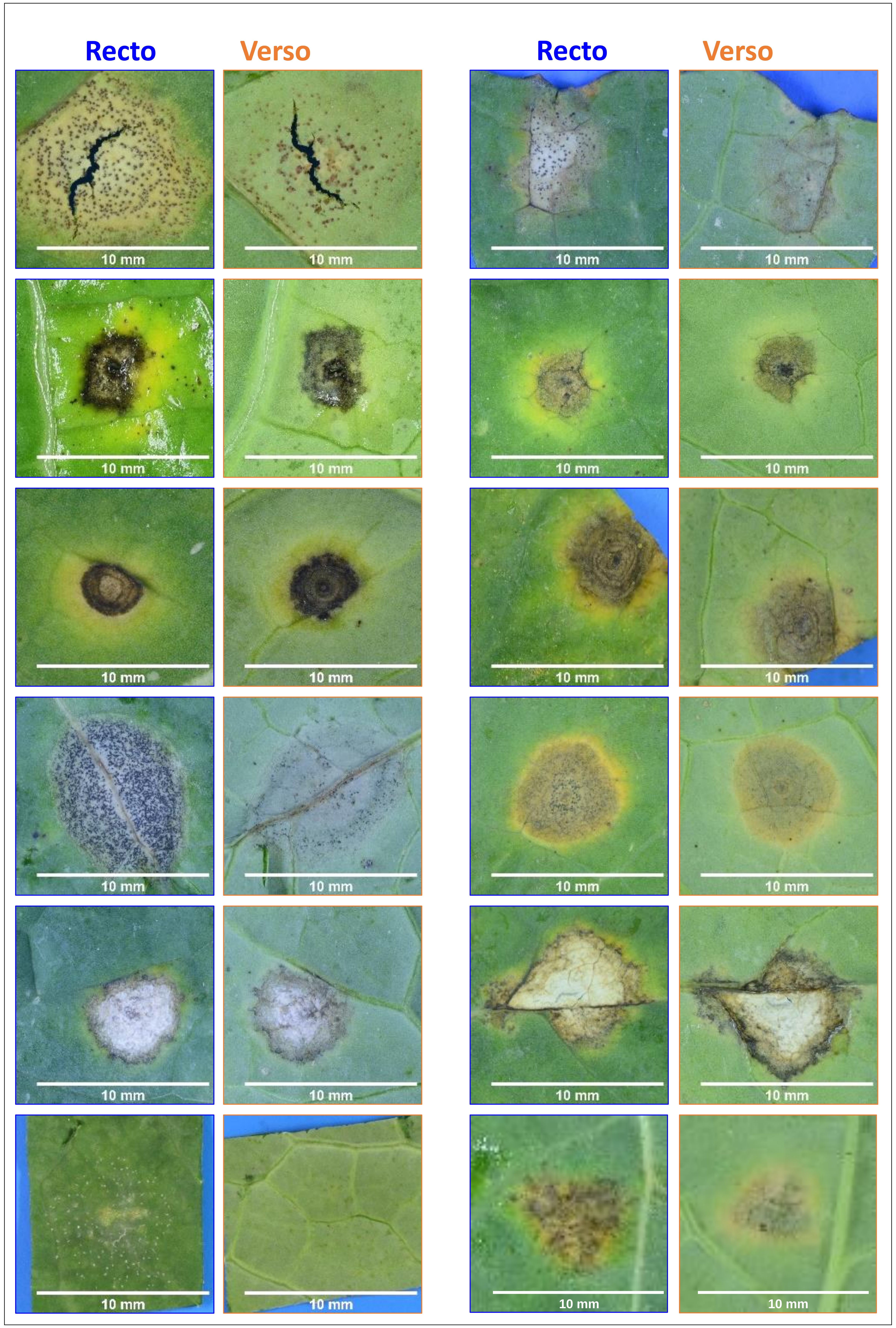


### Fig.S1

#### Distinctive criteria

**Phoma stem canker*, Leptosphaeria maculans***: Light grey leaf spot, often in a not fully regular shape. Black pycnidia visible to the naked eye, well distinct from one another. Diffuse border, generally without margin. Often the symptom stops at a leaf vein, which is black. Sometimes the leaf spot tears over: tissue collapse render them thinner. Verso of the symptom is green lighter than the leaf, and can bear some dots oosing pink spores (pycnidia)

**Phoma stem canker*, Leptosphaeria biglobosa***: Symptom seldom regular, spreading in all directions, generally stopped by leaf veins. Intense grey color, often with a darker dot in the middle (sometimes small tear). No or very few pycnida visible to the naked eye. Often the symptom is surrounded by a yellow halo. Verso of the symptom of intense grey color. Exist a larger and more regular form, with a black dot at the center. Sometimes dry necrosis of lighter grey. Exist a smaller form, black with a yellow halo.

**Black spot, *Alternaria brassicicola, A. brassicae***: Symptom of very regular shape, most often not stopping at leaf veins. Dark grey to blackish color. No pycnidia, and in most cases darker/lighter concentric circles are visible. There often is a yellow halo around the symptom. Verso of the symptom is dark grey to black.

**Ring spot, *Mycosphaerella brassicicola***: Symptom fully circular, most often not stopping at leaf veins. Dark grey color due to the many pycnidia, difficult to separate with the naked eye, smaller and more contiguous than these of *L. maculans*, sometimes arranged in circles. Often border slightly undulating and light green or yellow margin (not the same yellow than around *Alternaria* and *L. biglobosa*). Variable in color on the verso, but always slightly colored (beige) compared to *L. maculans*.

**White leaf spot, *Neopseudocercosporella capsellae***: Symptom of irregular shape, round or angular that sometimes stops at leaf margins. Whitish color, spot often with dark brown / black margin. Without visible pycnidia, but often marbling. On the verso, the presence of marbling/ dark veins is distinctive. Exist a more circular and white form.

***Light leaf spot, Pyrenopezziza brassicae***: Early stages of symptoms: Light grey discolored area, white sugar-like acervuli often forming a circular pattern around the discolored area. Either on the recto or the verso of the leaf. Late stages: dry beige necrosis of corky aspect and anarchic shape, with an intense yellow coloration (not the same yellow as the halo surrounding *Alternaria* or *L. biglobosa* symptoms). The old acervuli turn black and can be further around the crumpled symptom. In severe forms, the leaf can be crinkled and symptoms cracked.

### Fig.S2 Phoma stem canker, *Leptosphaeria maculans*

**Phoma stem canker, *Leptosphaeria maculans***: Light grey leaf spot, often in a not fully regular shape. Black pycnidia visible to the naked eye, well distinct from one another. Diffuse border, generally without margin. Often the symptom stops at a leaf vein, which is black. Sometimes the leaf spot tears over: tissue collapse render them thinner. Verso of the symptom is green lighter than the leaf, and can bear some dots oosing pink spores (pycnidia)

**2a. Typical symptoms**: exemples respecting most of the distinctive criteria for the species

Recto

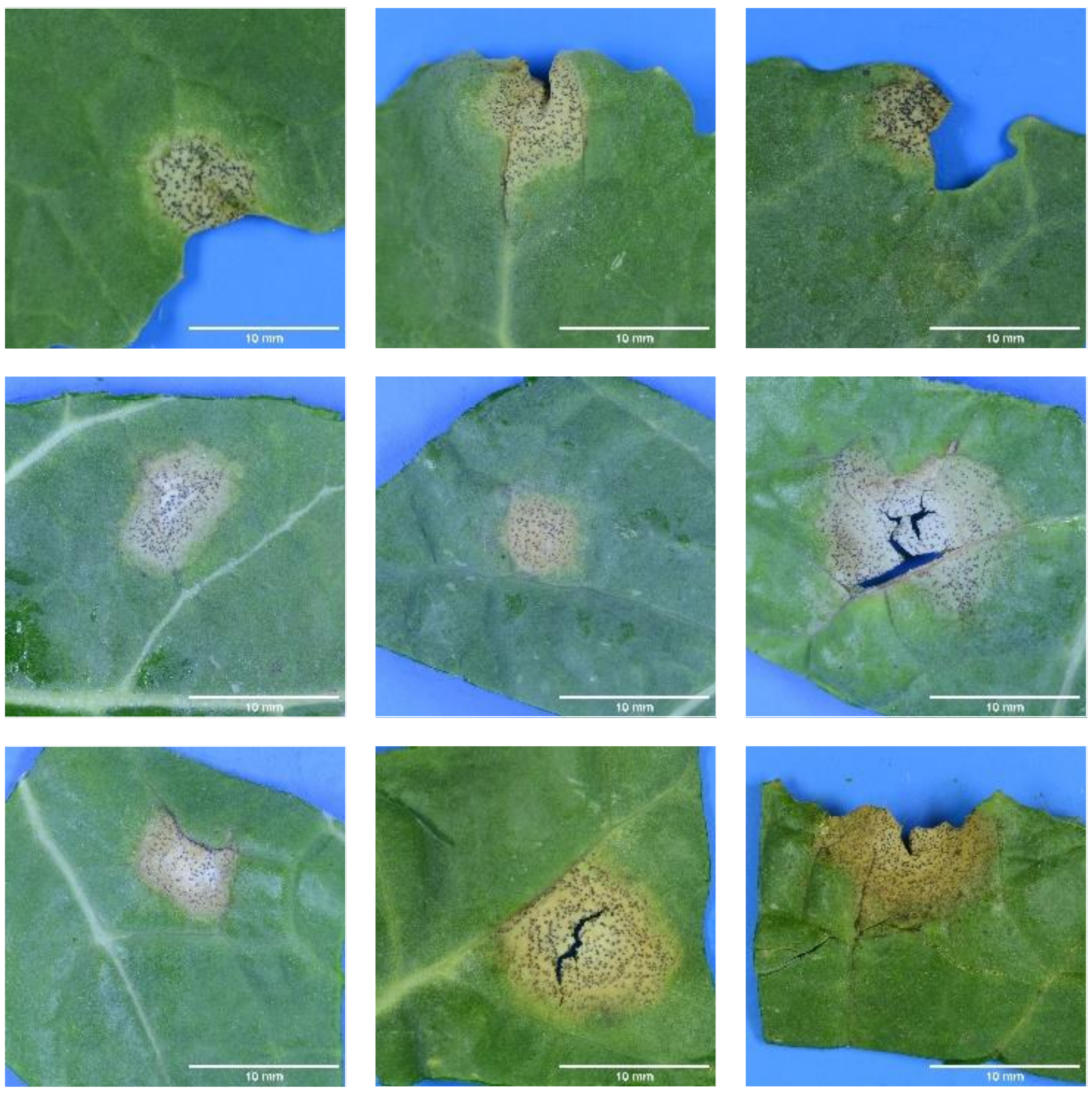

Verso

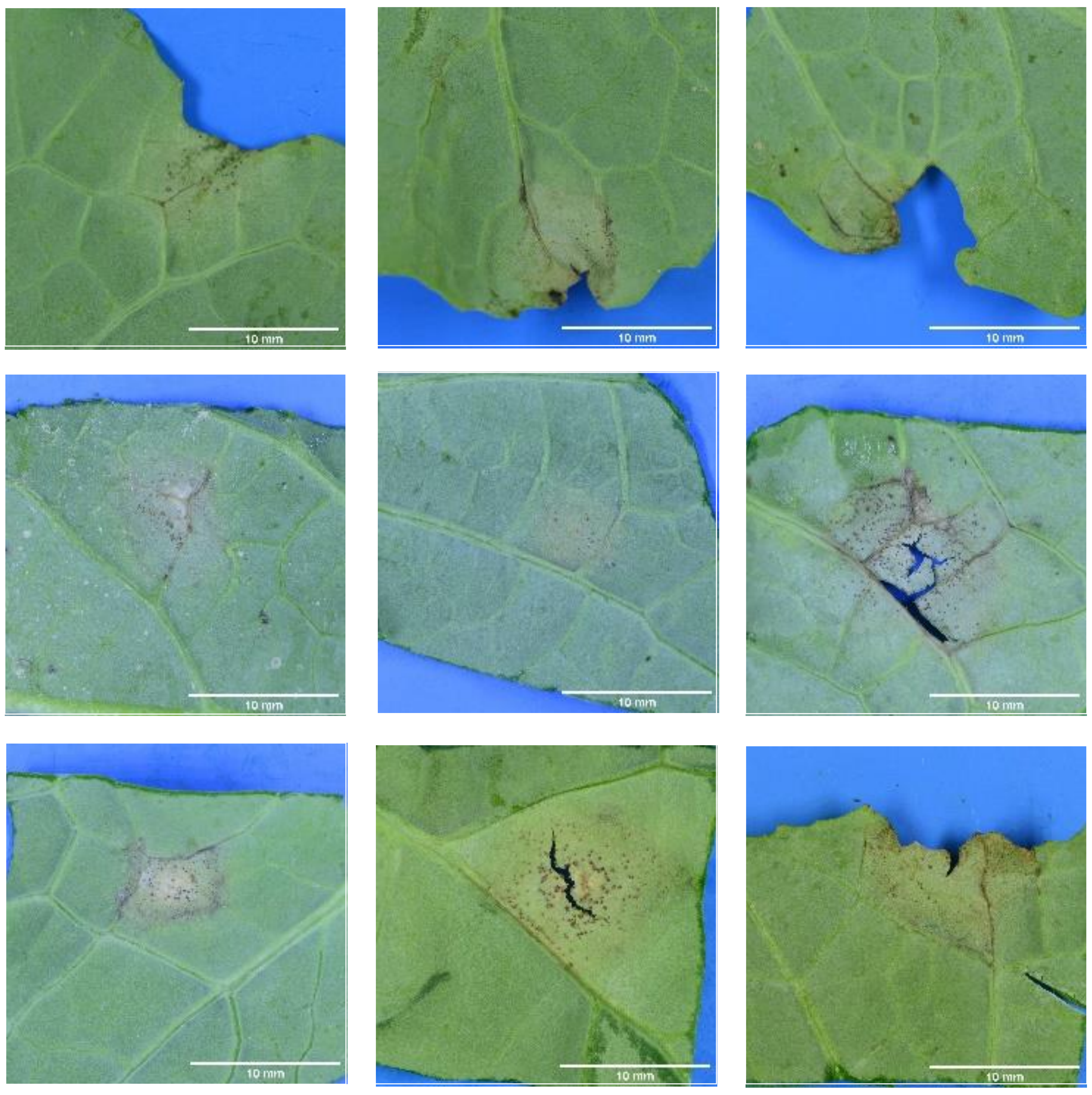

**2b. Atypical symptoms:** examples of variations respecting only some of the distinctive criteria

Recto

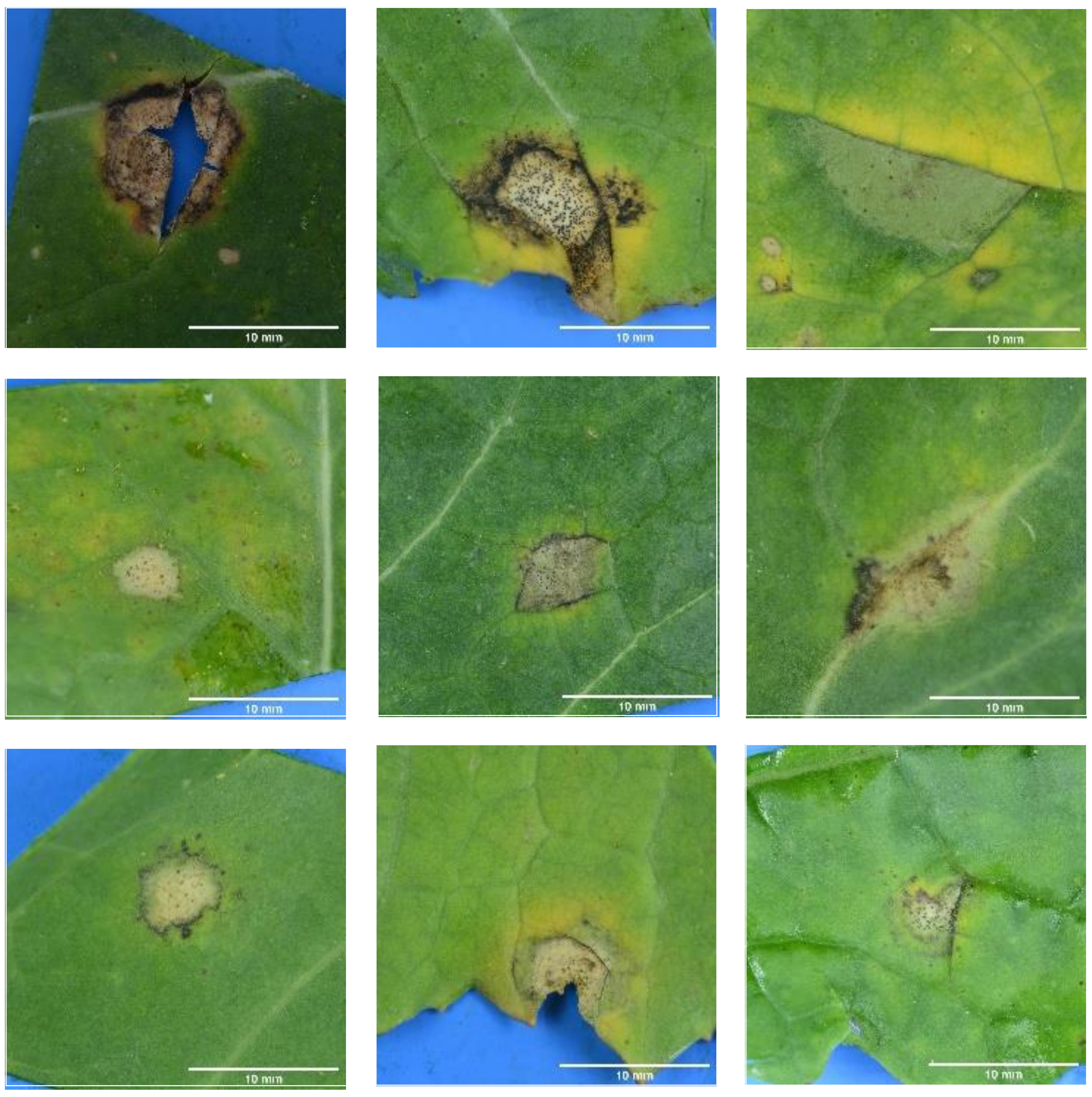

Verso

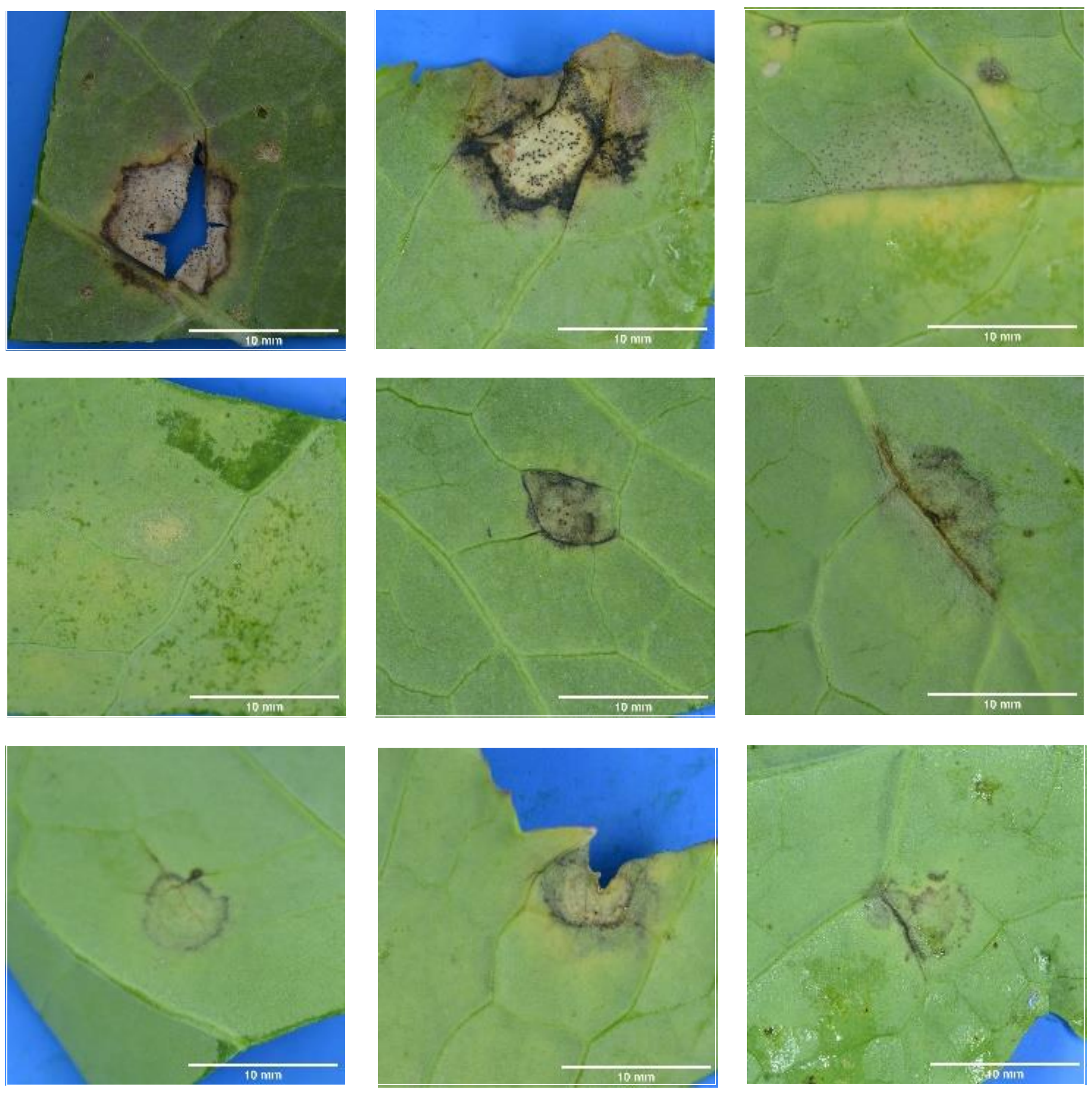

### Fig.S3 Phoma stem canker, *Leptosphaeria biglobosa*

**Phoma stem canker, *Leptosphaeria biglobosa***: Symptom seldom regular, spreading in all directions, generally stopped by leaf veins. Intense grey color, often with a darker dot in the middle (sometimes small tear). No or very few pycnida visible to the naked eye. Often the symptom is surrounded by a yellow halo. Verso of the symptom of intense grey color. Exist a larger and more regular form, with a black dot at the center. Sometimes dry necrosis of lighter grey. Exist a smaller form, black with a yellow halo.

**3a. Typical symptoms**: exemples respecting most of the distinctive criteria for the species

Recto

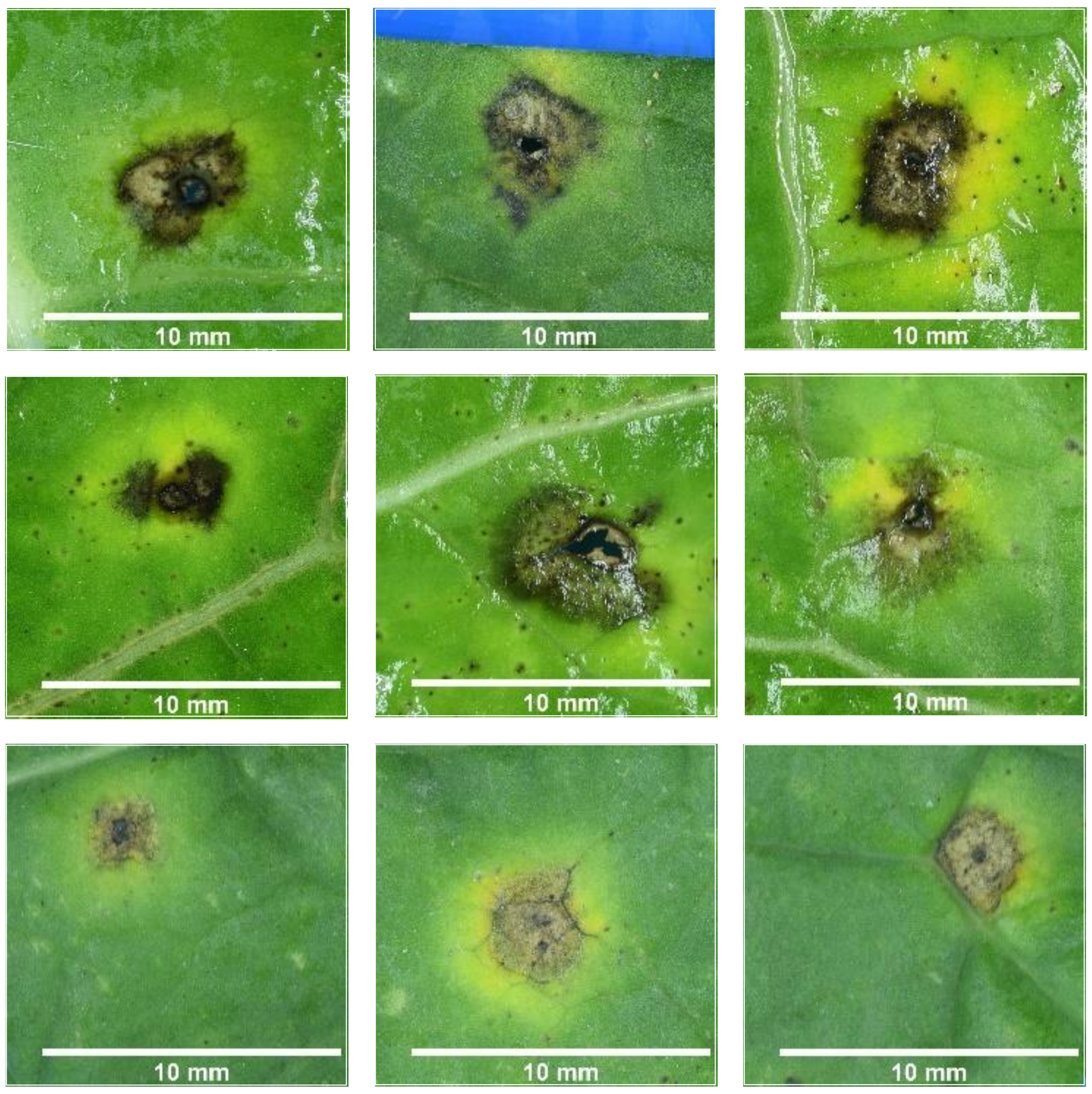


Verso

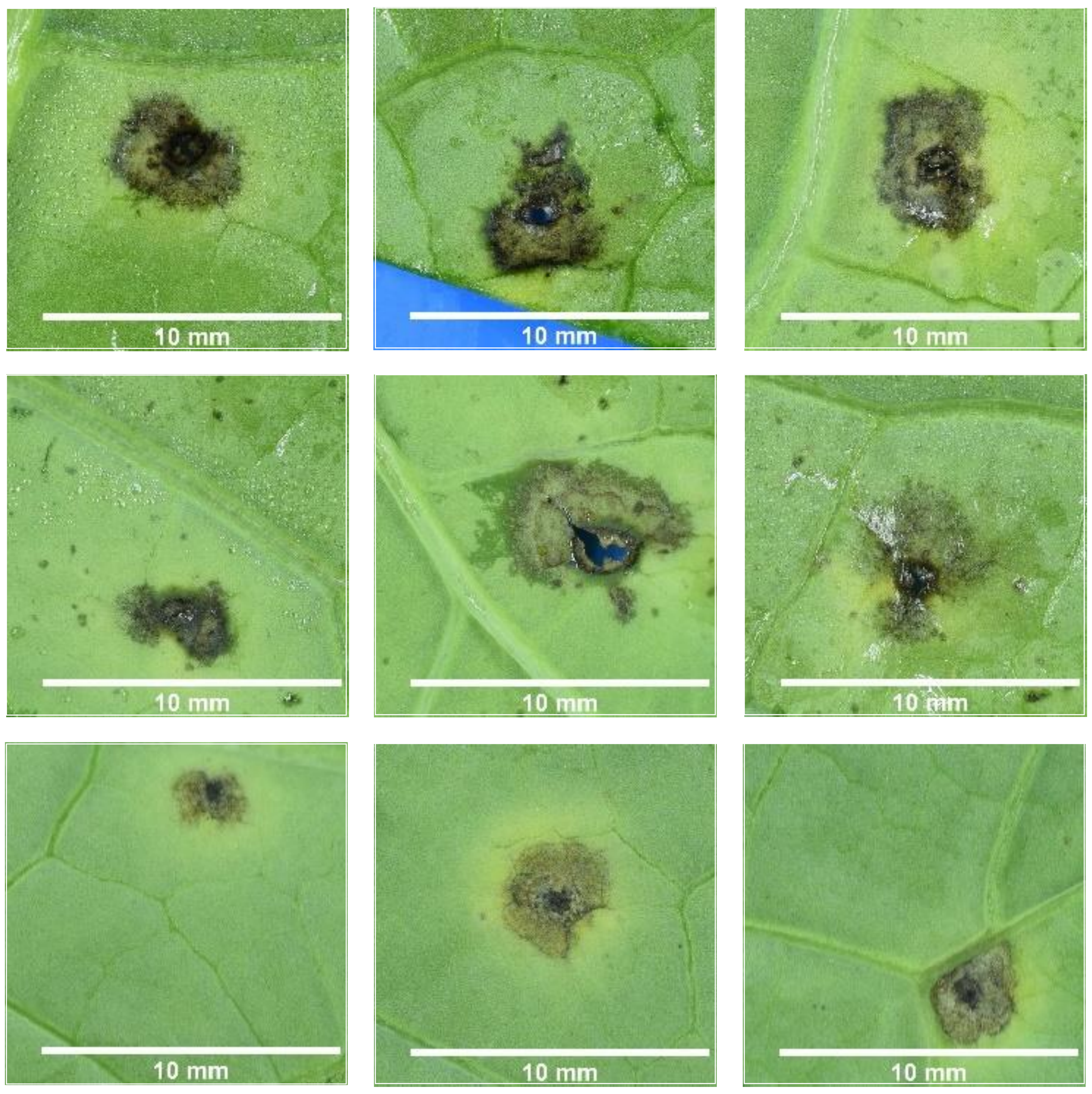


**3b. Atypical symptoms:** examples of variations respecting only some of the distinctive criteria

Recto

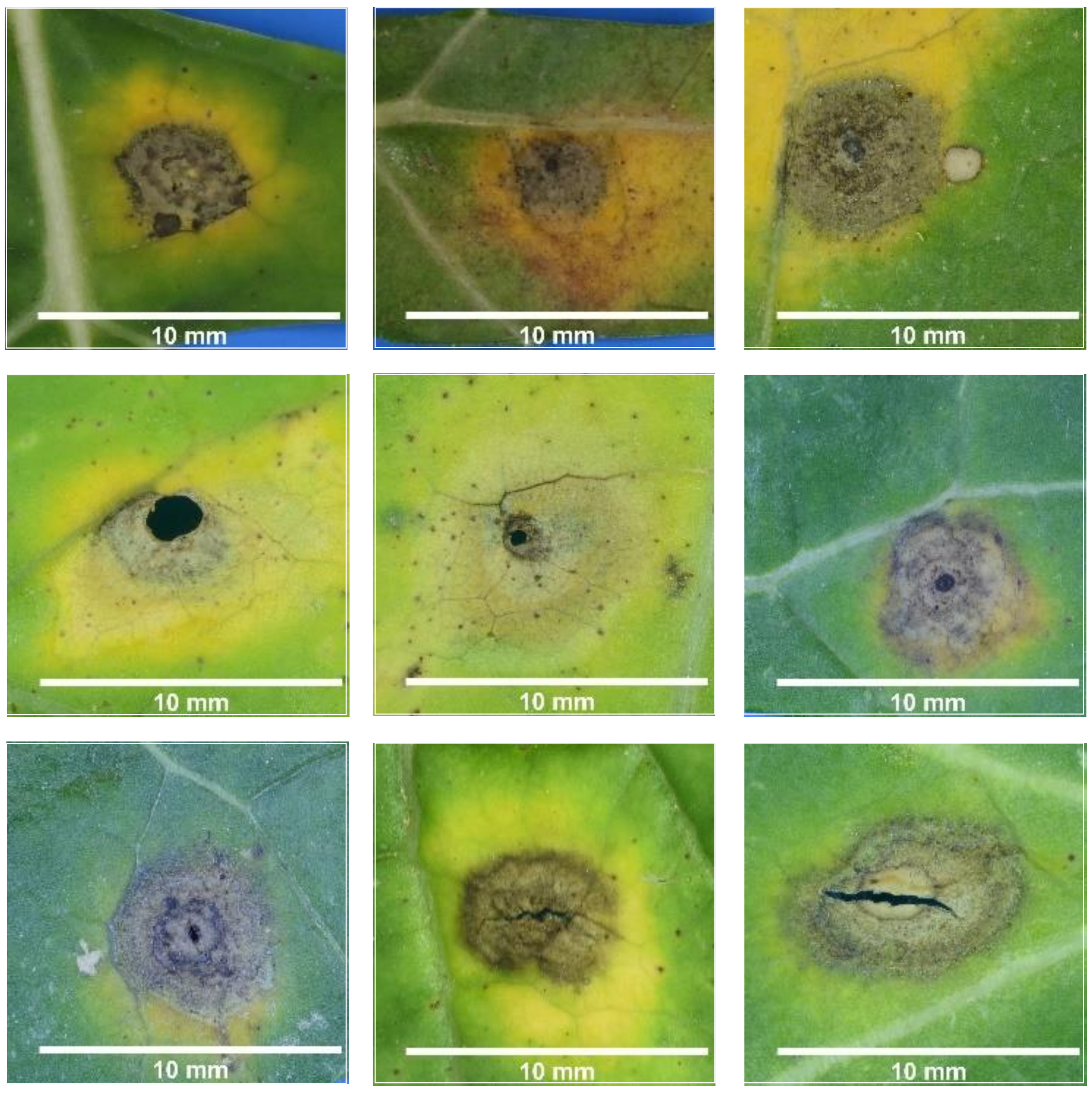


Verso

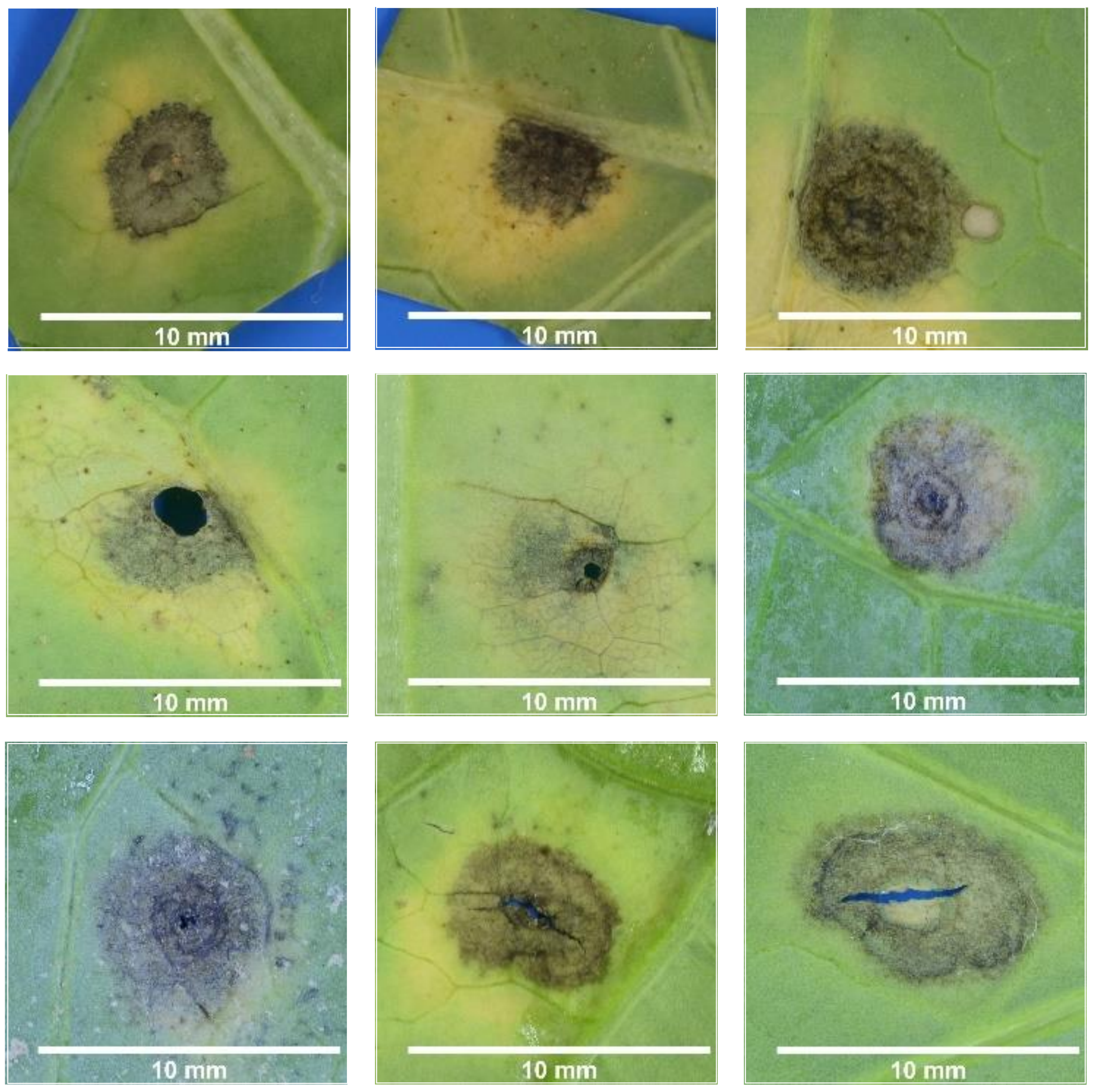

### Fig.S4 White spot, *Neopseudocercosporella capsellae*

**White leaf spot, *Neopseudocercosporella capsellae***: Symptom of irregular shape, round or angular that sometimes stops at leaf margins. Whitish color, spot often with dark brown / black margin. Without visible pycnidia, but often marbling. On the verso, the presence of marbling/ dark veins is distinctive. Exist a more circular and white form

**4a. Typical symptoms**: exemples respecting most of the distinctive criteria for the species

Recto

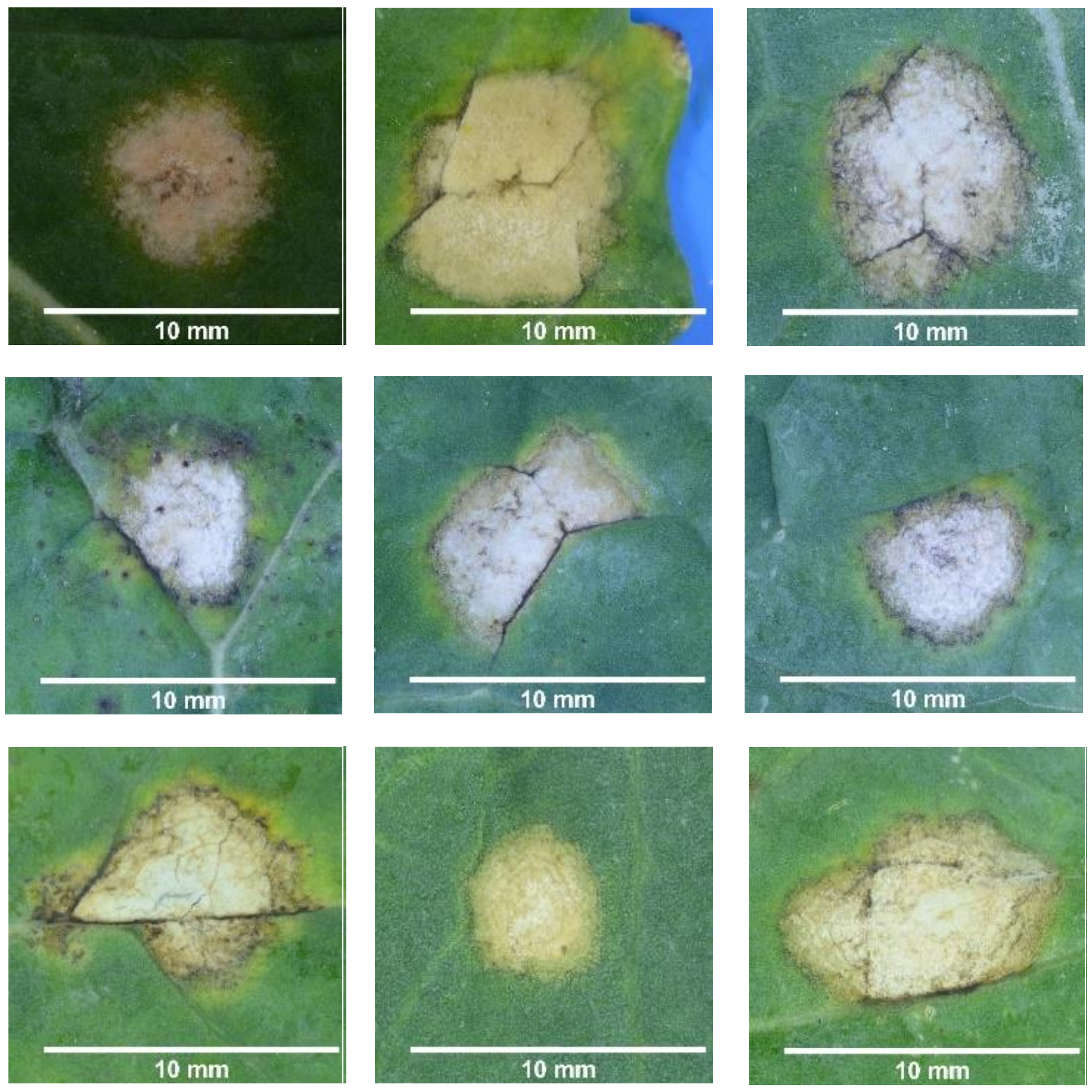


Verso

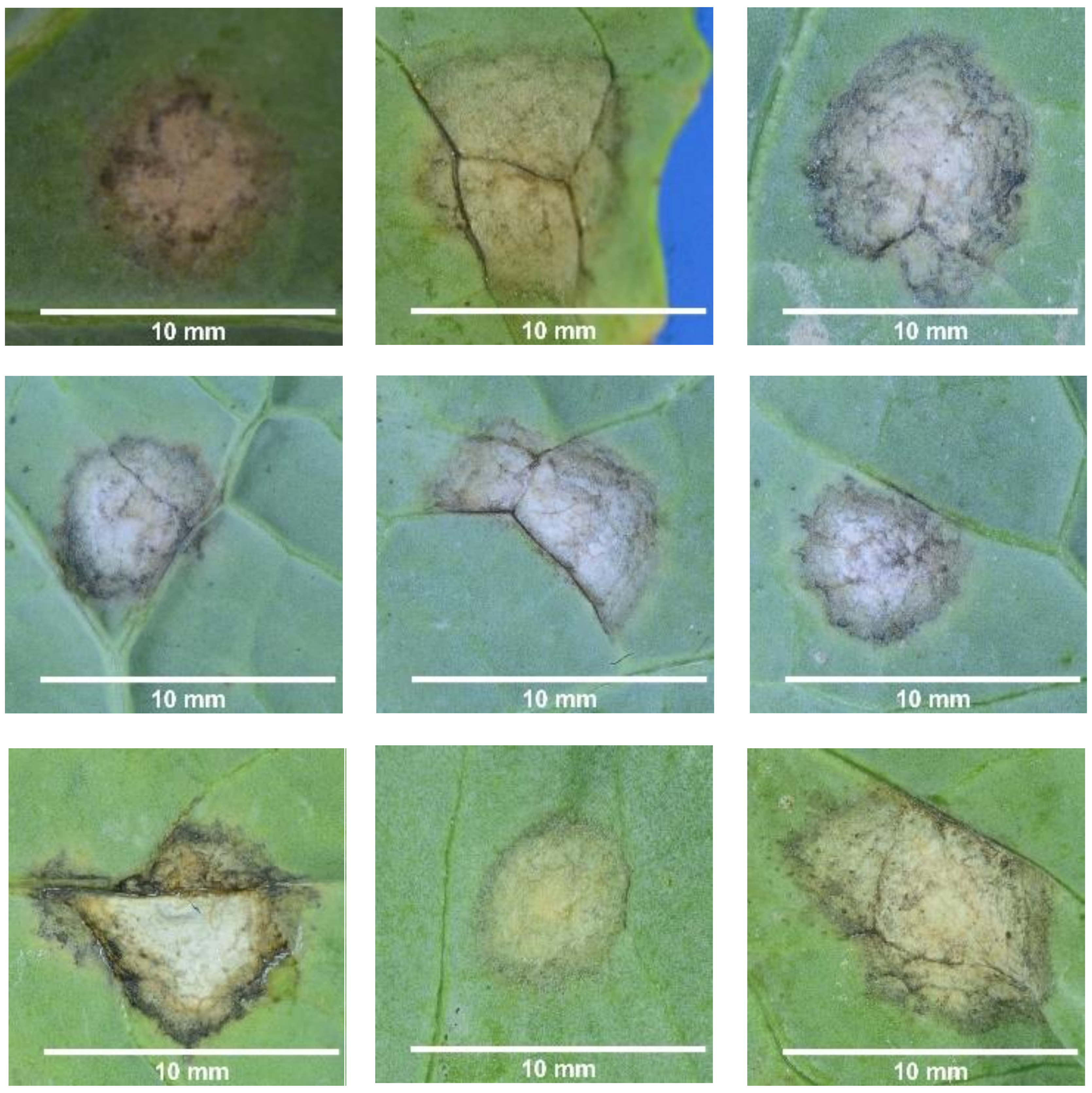


**4b. Atypical symptoms:** examples of variations respecting only some of the distinctive criteria

Recto

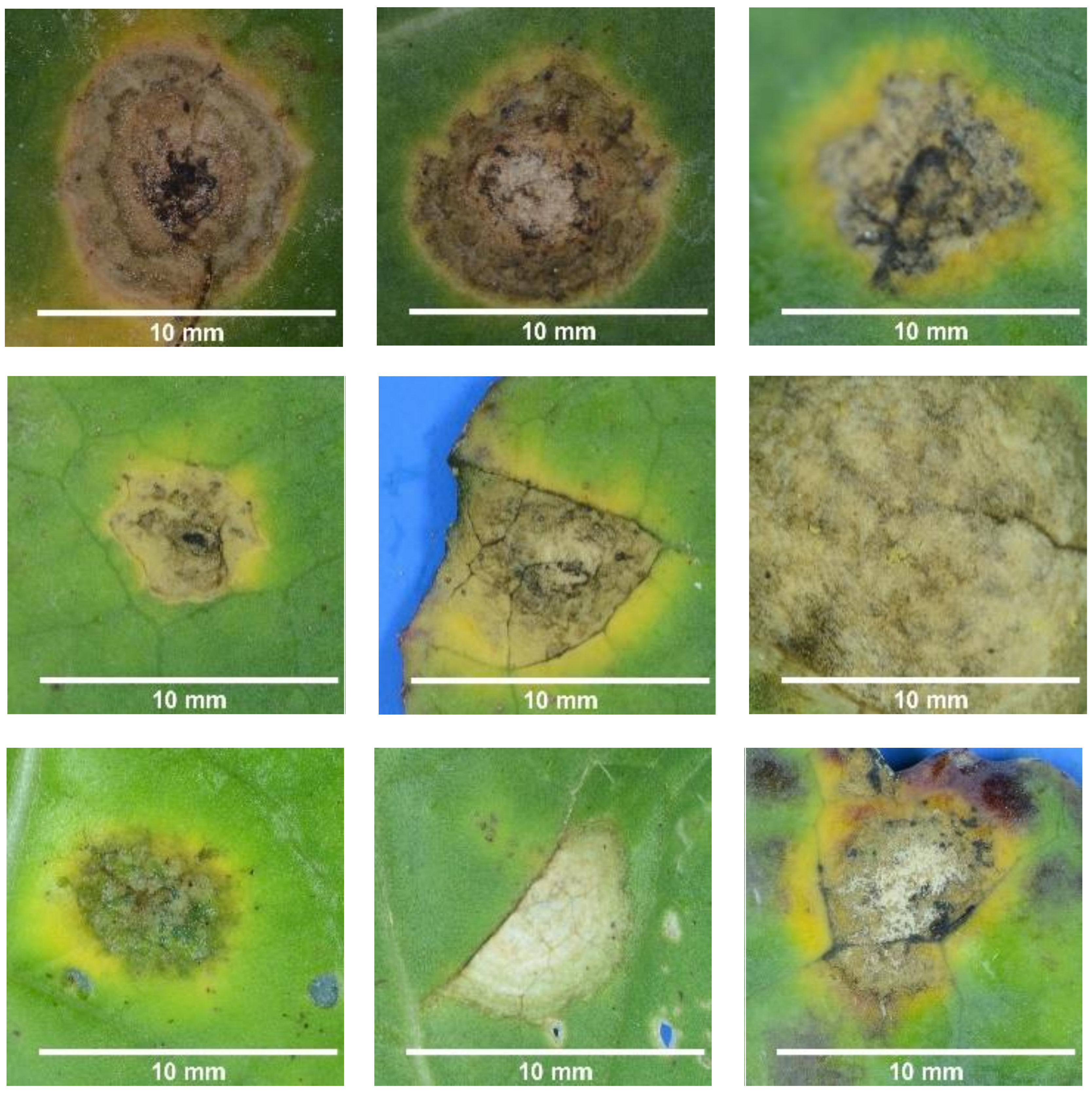


Verso

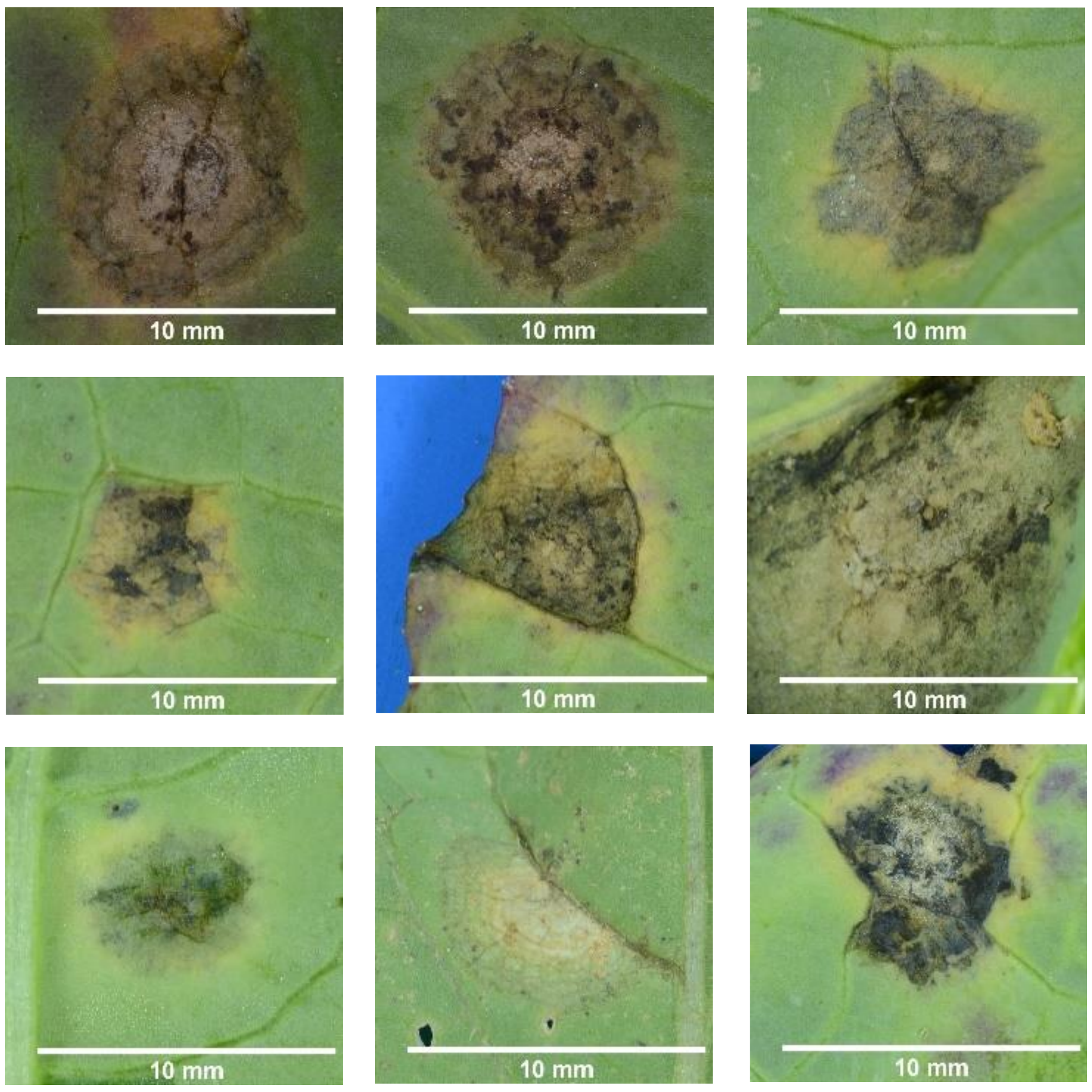

### Fig.S5 Ring spot, *Mycosphaerella brassicicola*

**Ring spot, *Mycosphaerella brassicicola***: Symptom fully circular, most often not stopping at leaf veins. Dark grey color due to the many pycnidia, difficult to separate with the naked eye, smaller and more contiguous than these of *L. maculans*, sometimes arranged in circles. Often border slightly undulating and light green or yellow margin (not the same yellow than around *Alternaria* and *L. biglobosa*). Variable in color on the verso, but always slightly colored (beige) compared to *L. maculans*.

**5a. Typical symptoms**: exemples respecting most of the distinctive criteria for the species

Recto

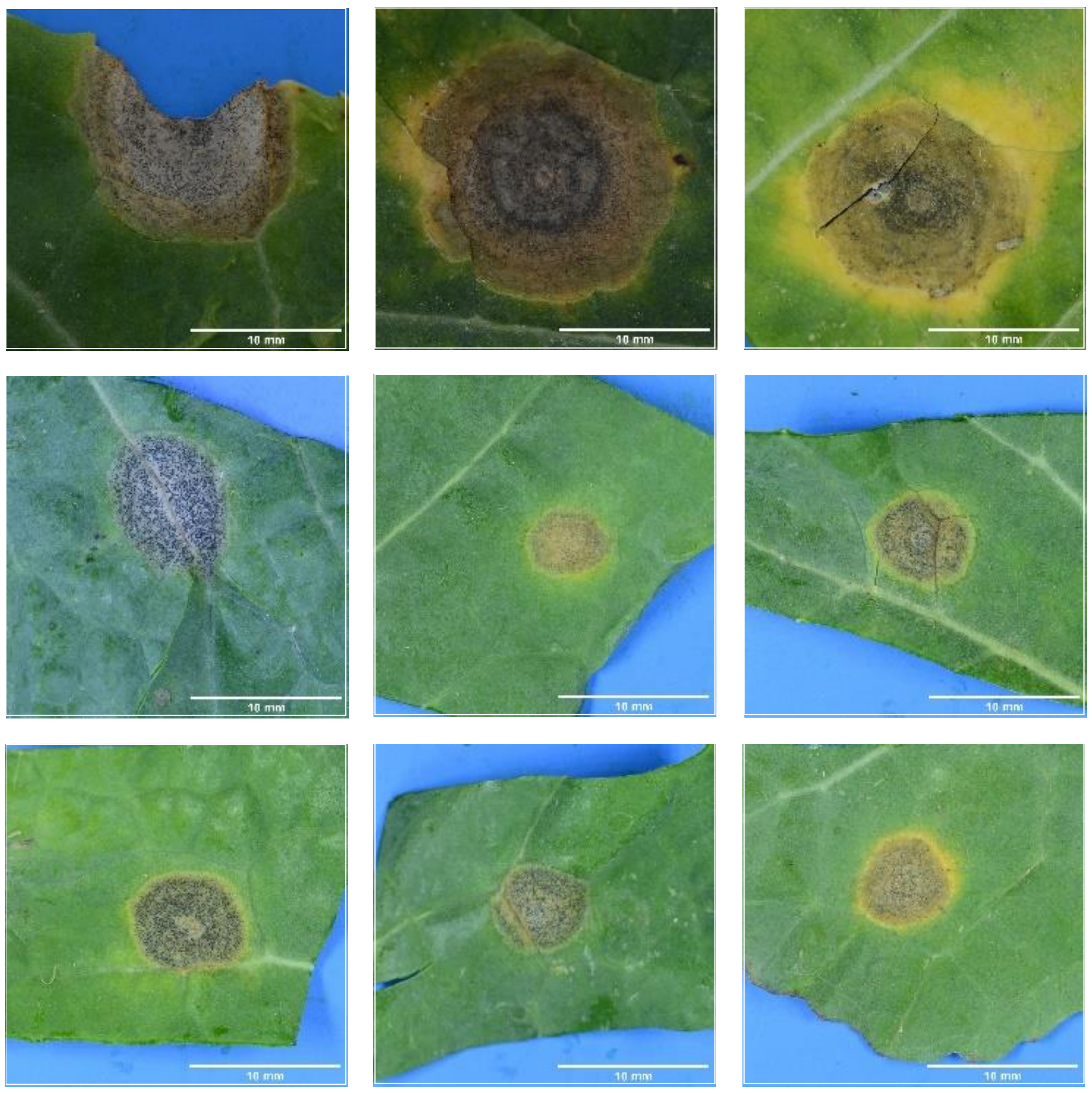

Verso

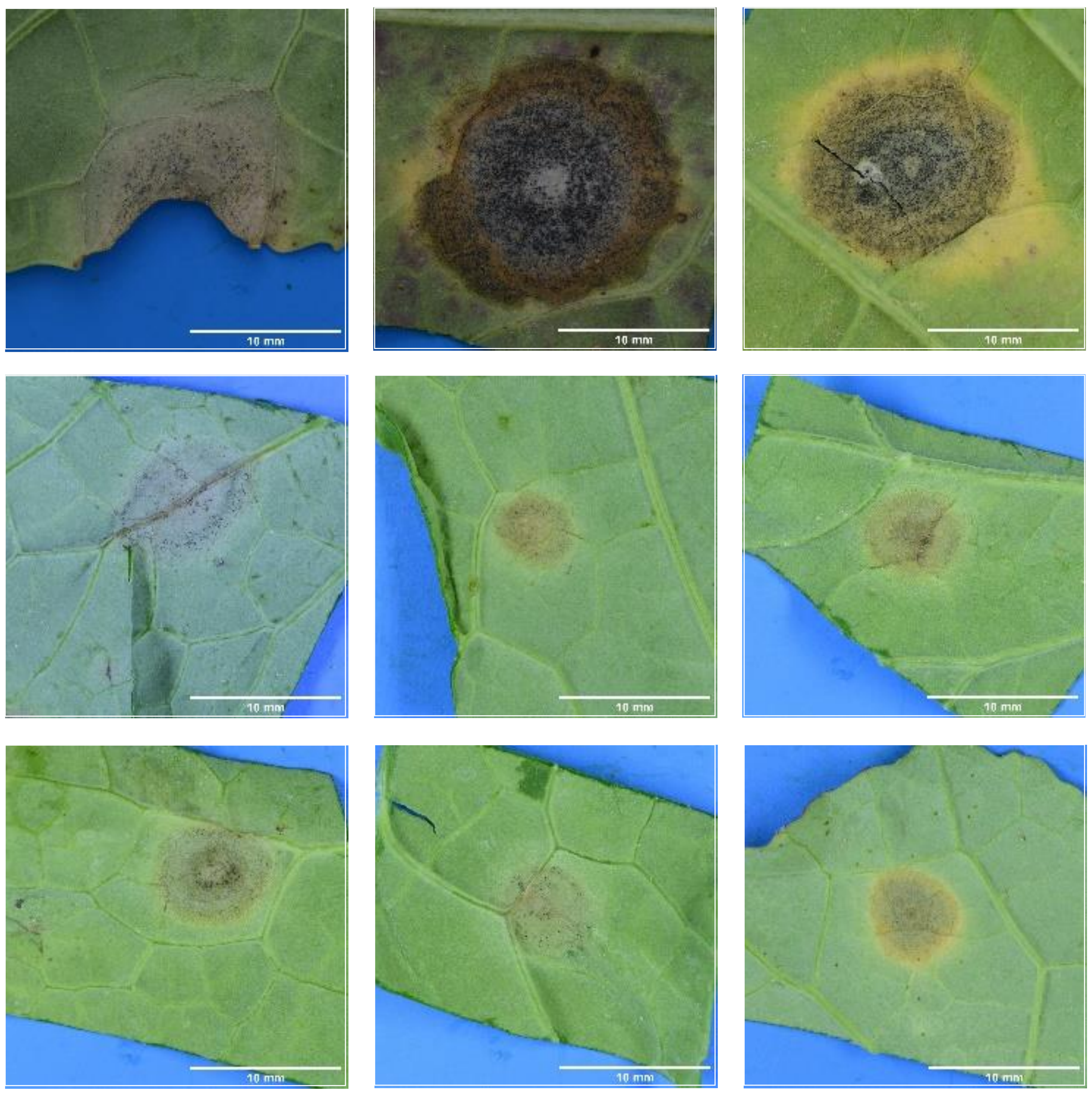

**5b. Atypical symptoms:** examples of variations respecting only some of the distinctive criteria

Recto

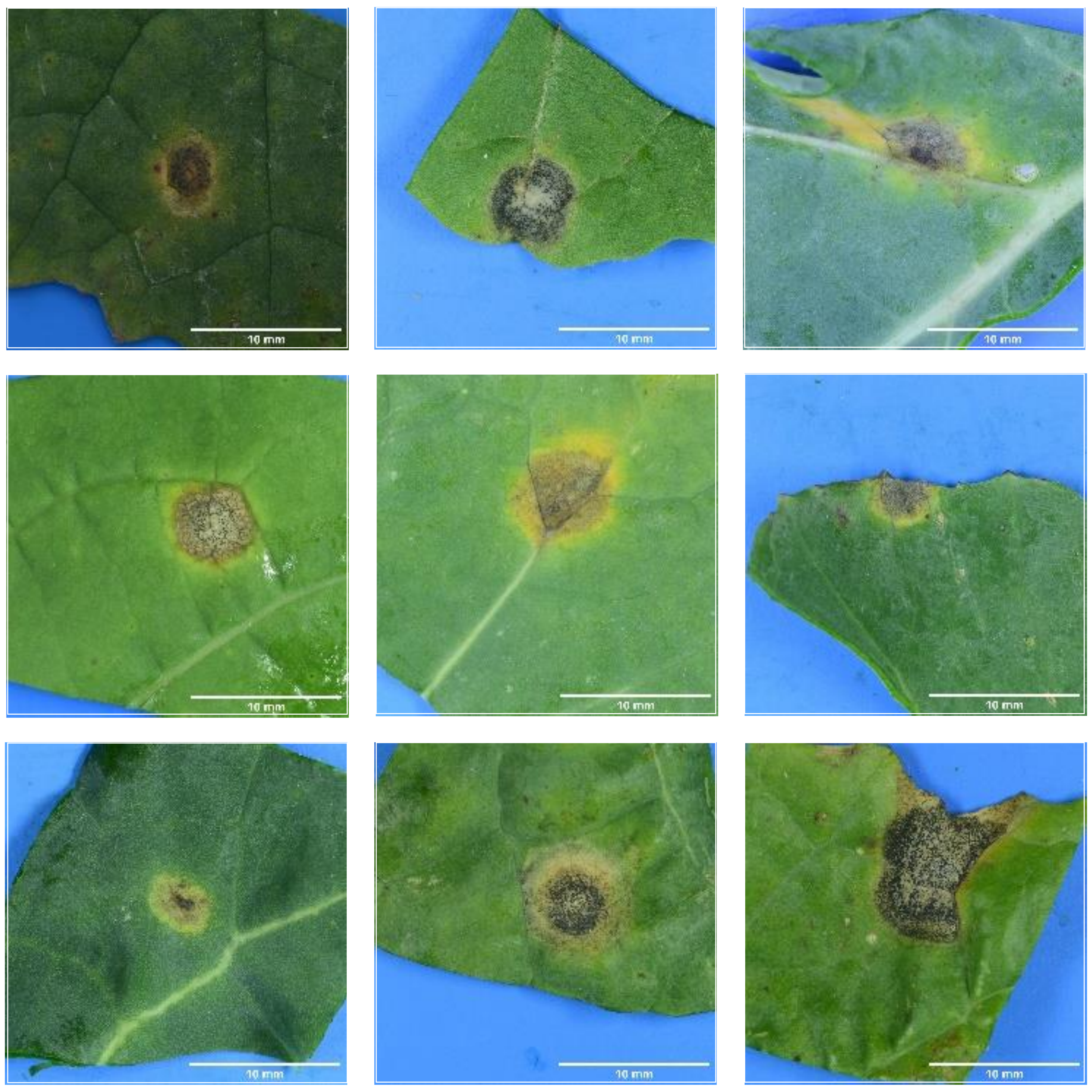

Verso

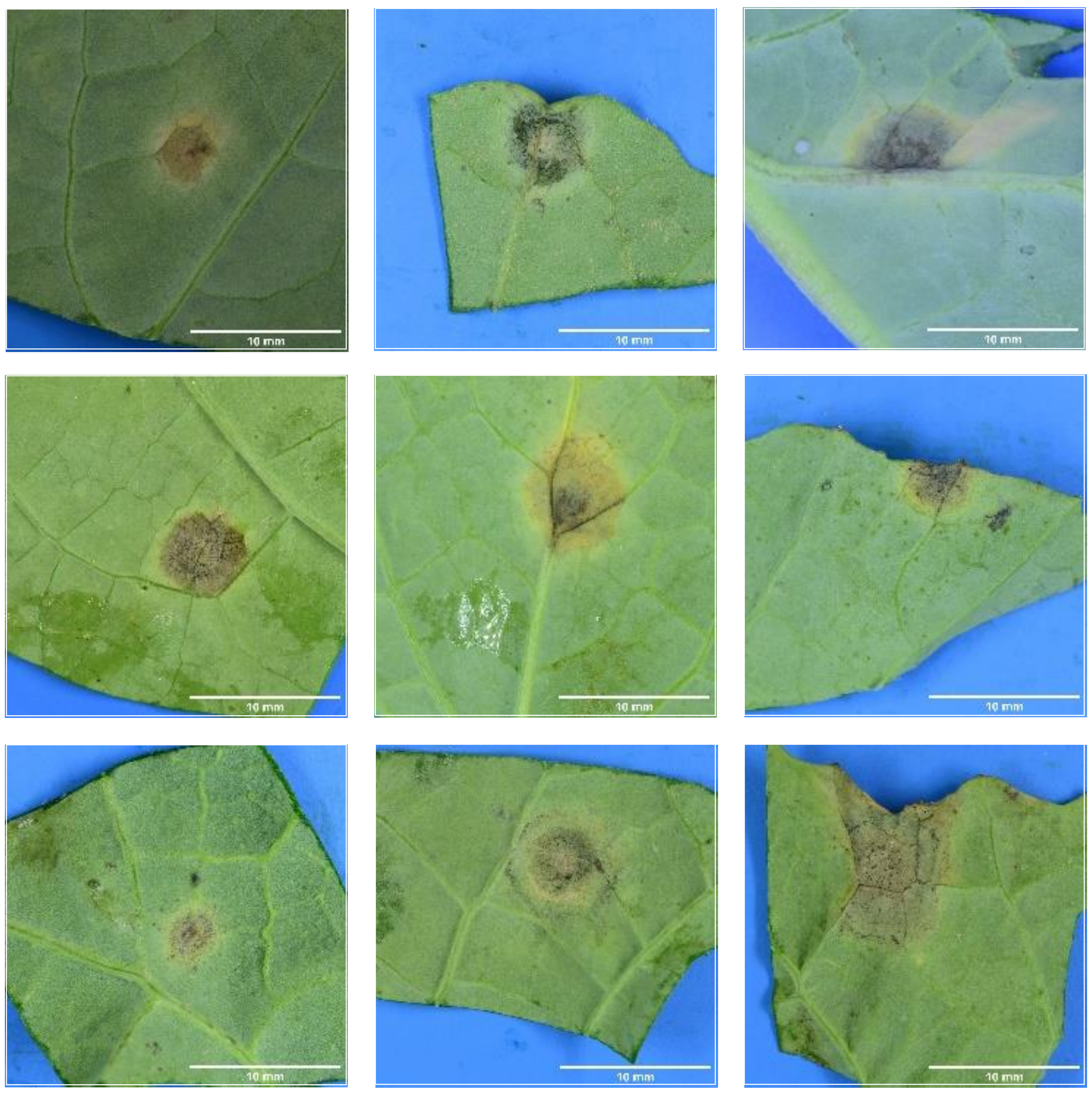

### Fig.S6 Black spot, *Alternaria brassicicola, A. brassicae*

**Black spot, *Alternaria brassicicola, A. brassicae***: Symptom of very regular shape, most often not stopping at leaf veins. Dark grey to blackish color. No pycnidia, and in most cases darker/lighter concentric circles are visible. There often is a yellow halo around the symptom. Verso of the symptom is dark grey to black.

**6a. Typical symptoms**: exemples respecting most of the distinctive criteria for the species

Recto

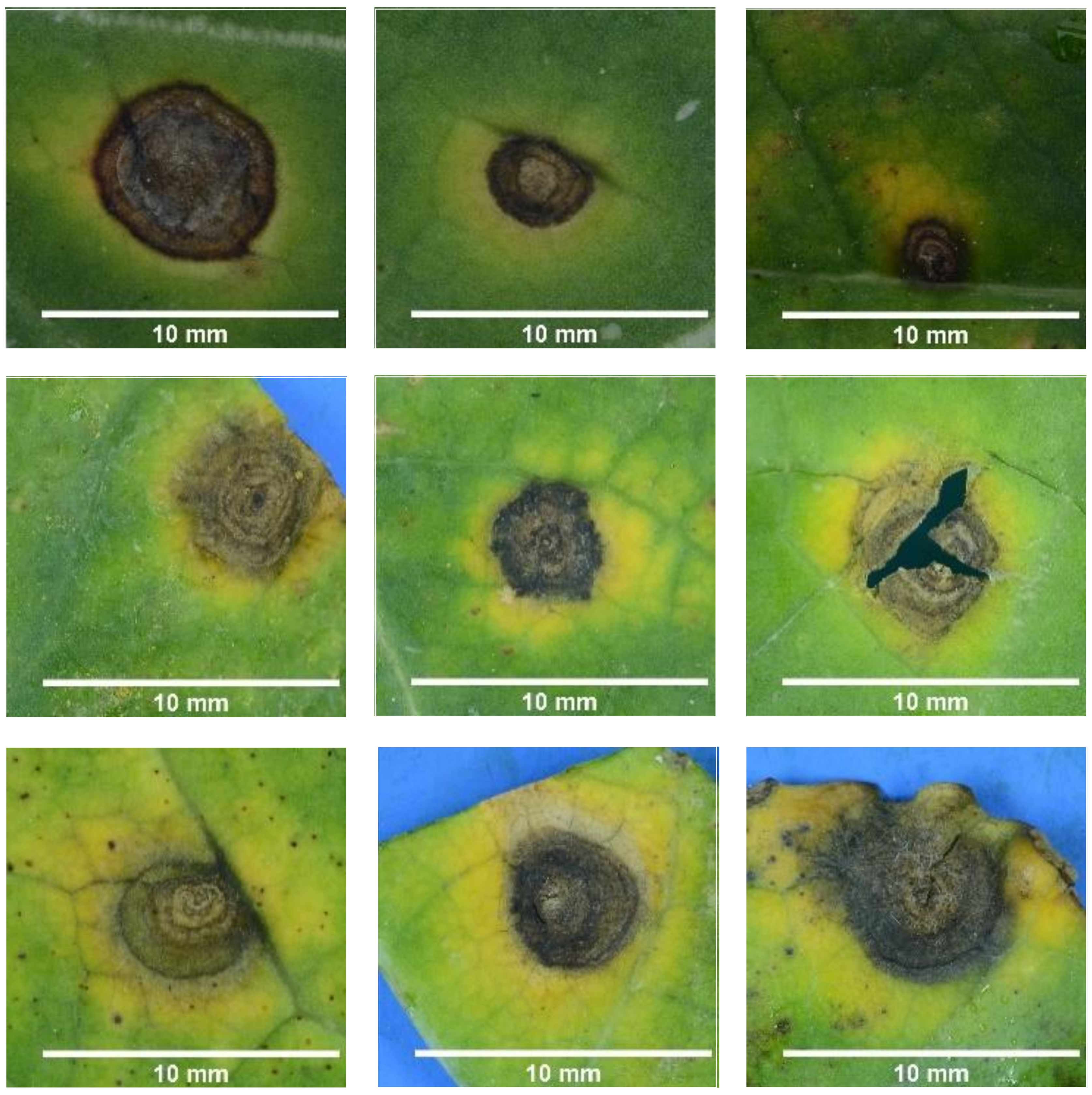


Verso

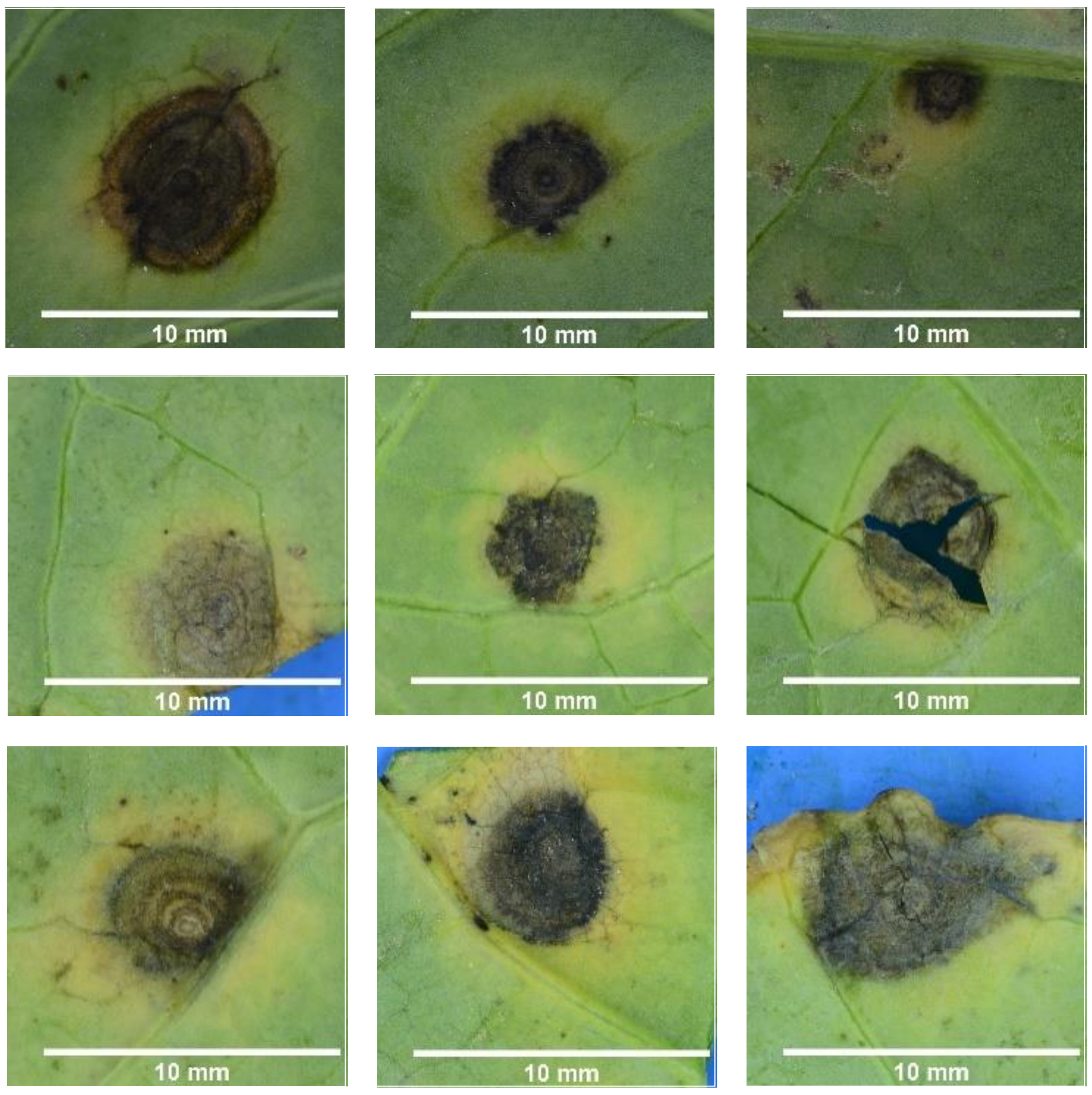


**6b. Atypical symptoms:** examples of variations respecting only some of the distinctive criteria

Recto

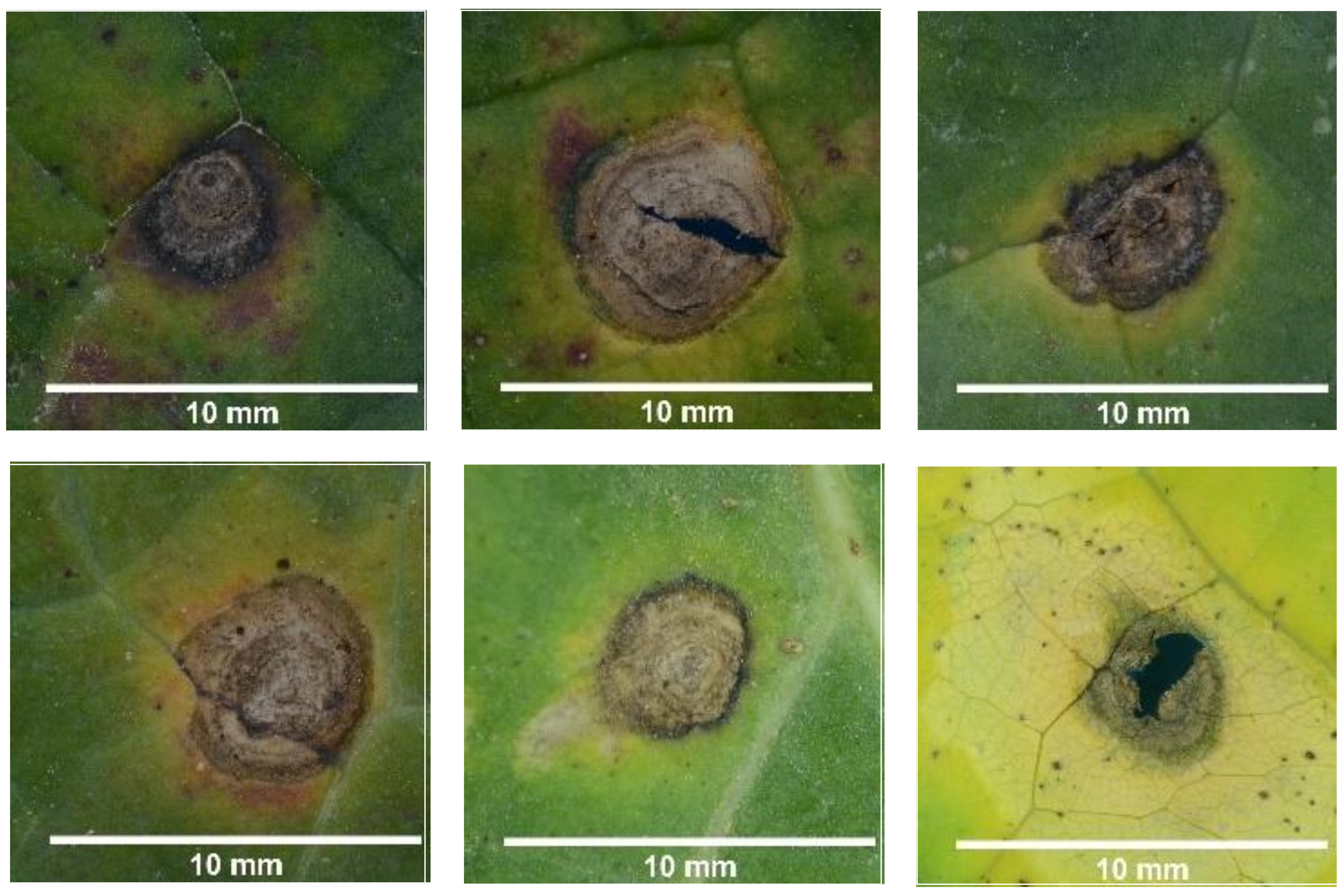


Verso

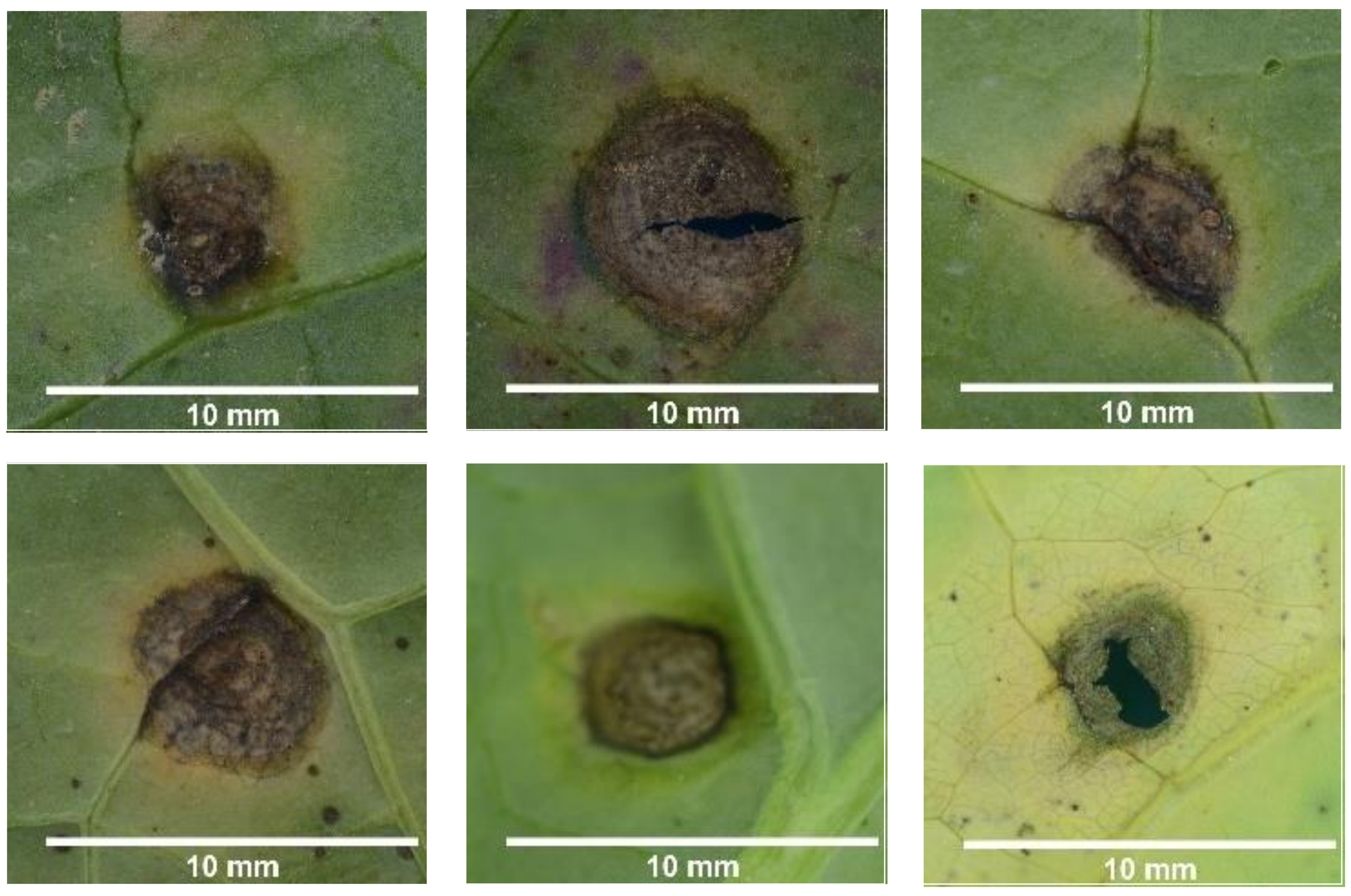

### Fig.S7 Light leaf spot, *Pyrenopezziza brassicae* :

***Light leaf spot, Pyrenopezziza brassicae***: *Early stages* of symptoms: Light grey discolored area, white sugar-like acervuli often forming a circular pattern around the discolored area. Either on the recto or the verso of the leaf. *Late stages:* dry beige necrosis of corky aspect and anarchic shape, with an intense yellow coloration (not the same yellow as the halo surrounding *Alternaria* or *L. biglobosa* symptoms). The old acervuli turn black and can be further around the crumpled symptom. In severe forms, the leaf can be crinkled and symptoms cracked.

#### 7a. Typical symptoms: exemples respecting most of the distinctive criteria for the species

Recto

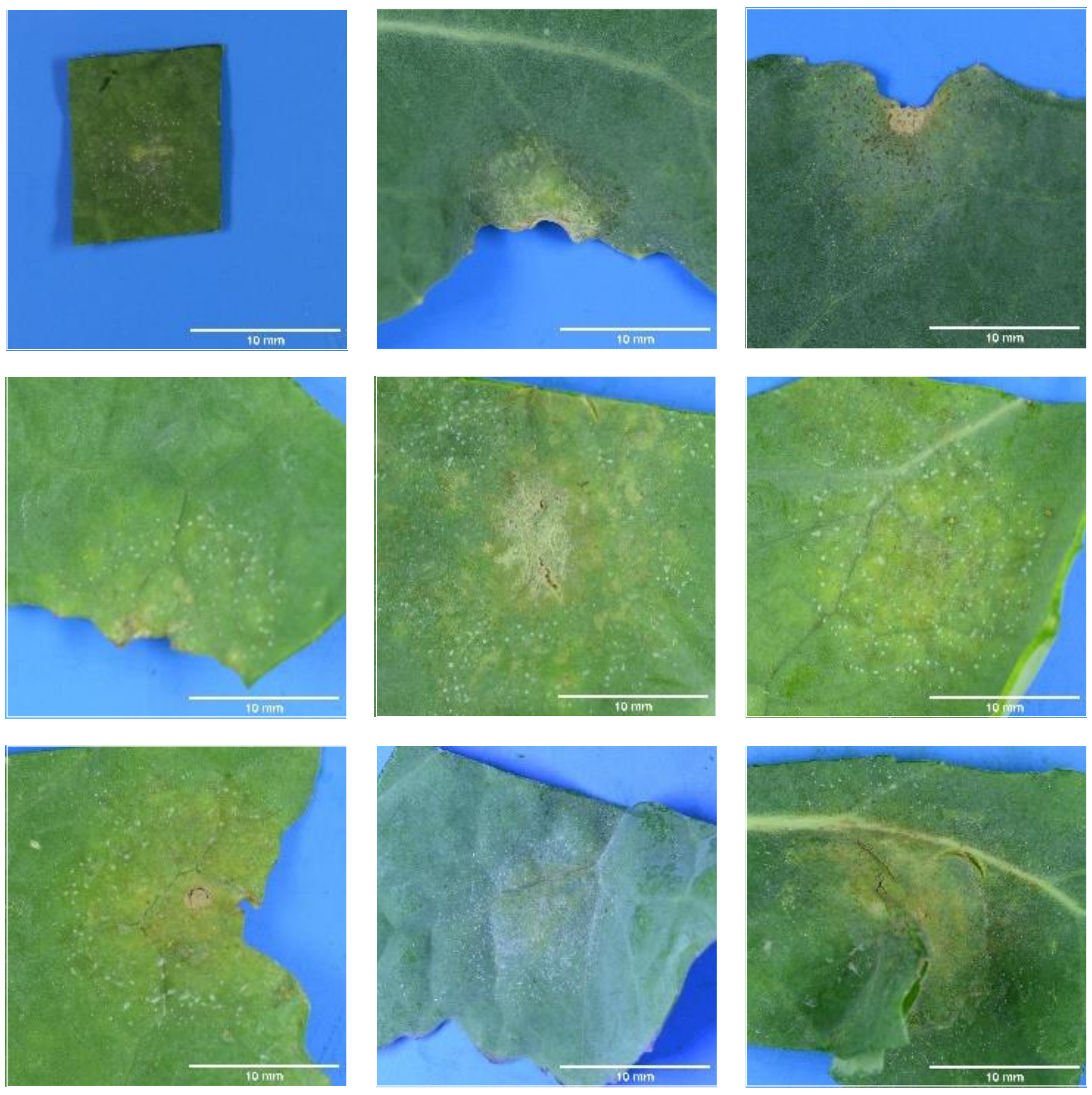

Verso

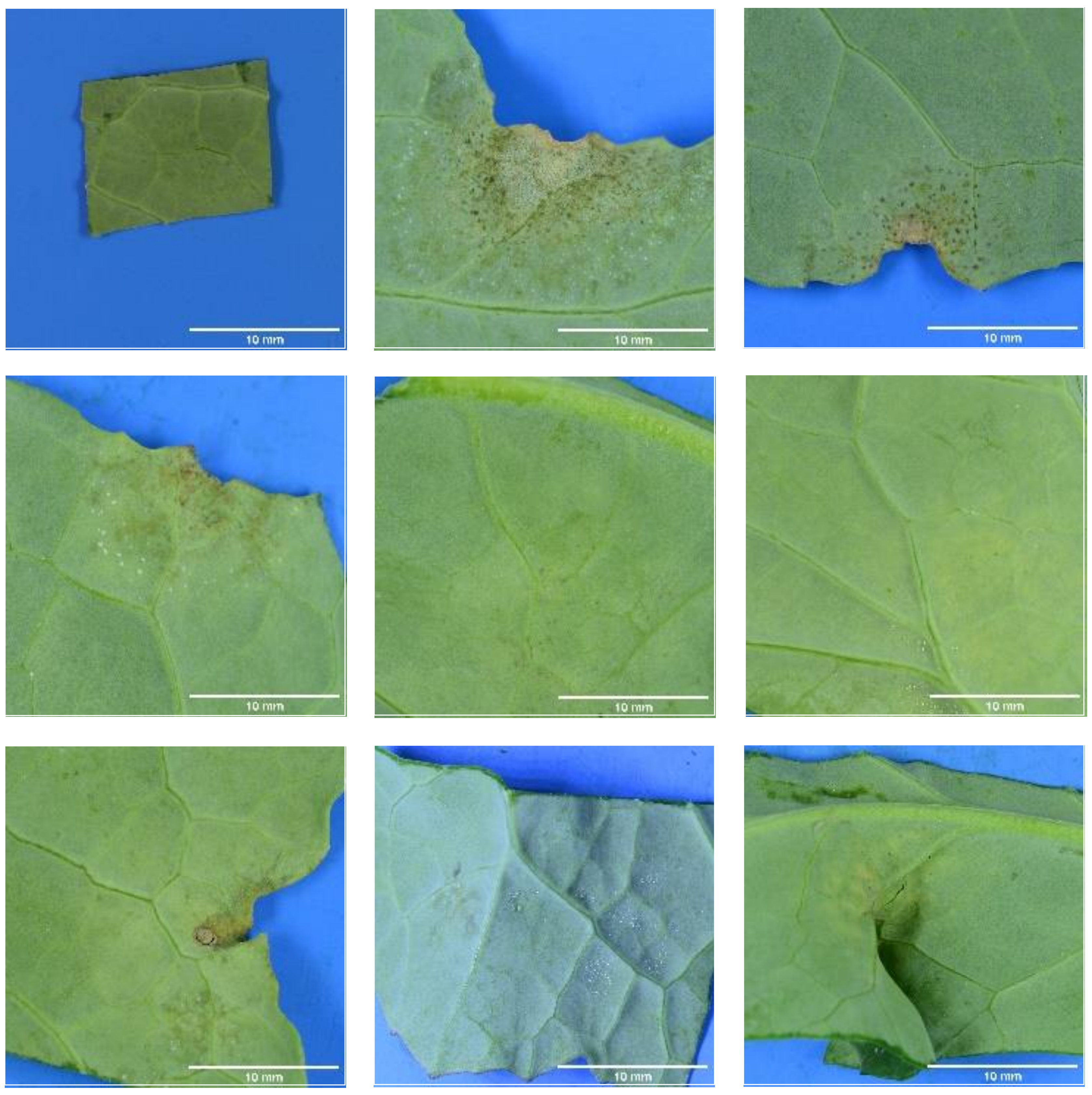

#### 7b. Atypical symptoms: examples of variations respecting only some of the distinctive criteria

Recto

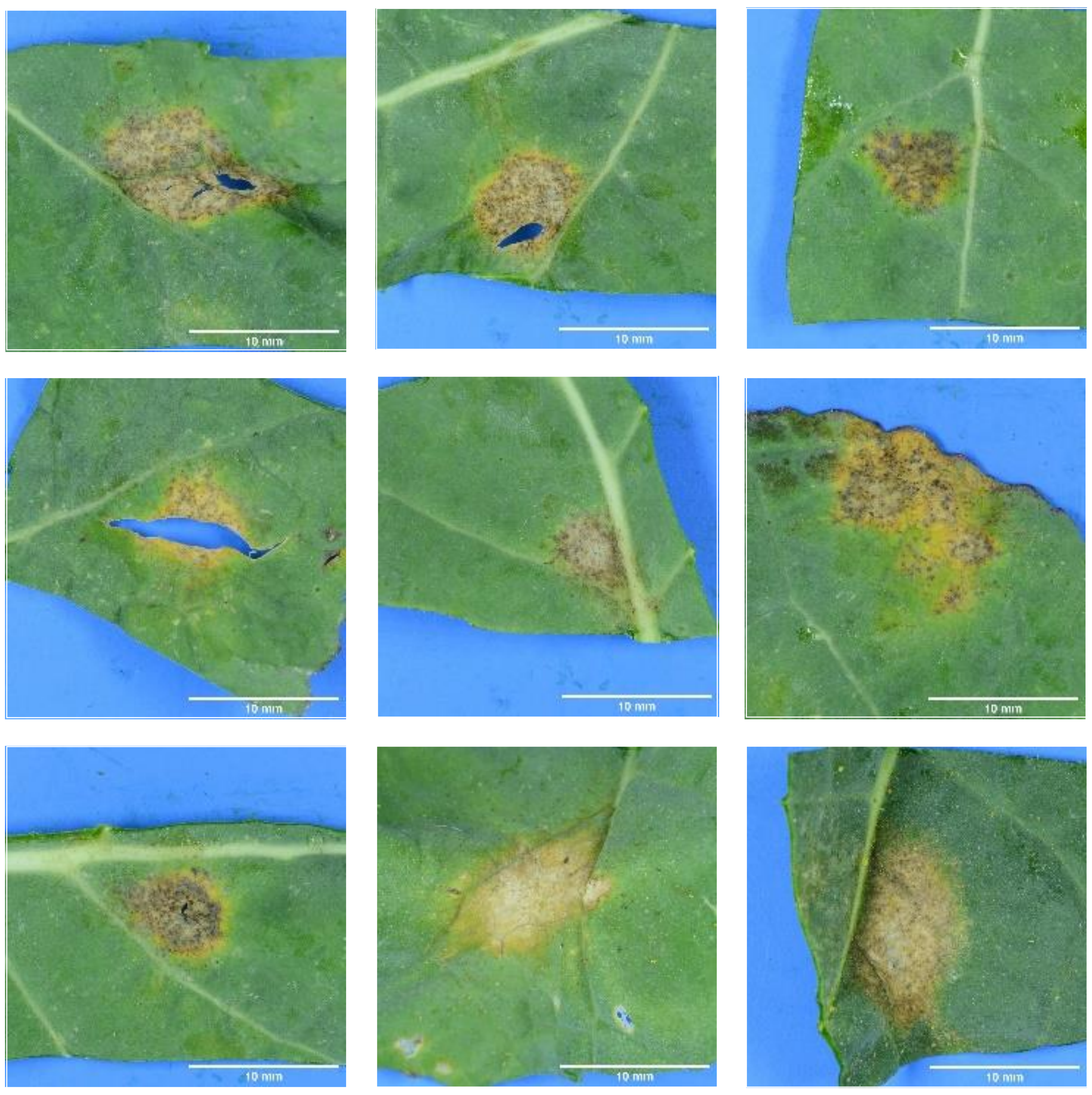

Verso

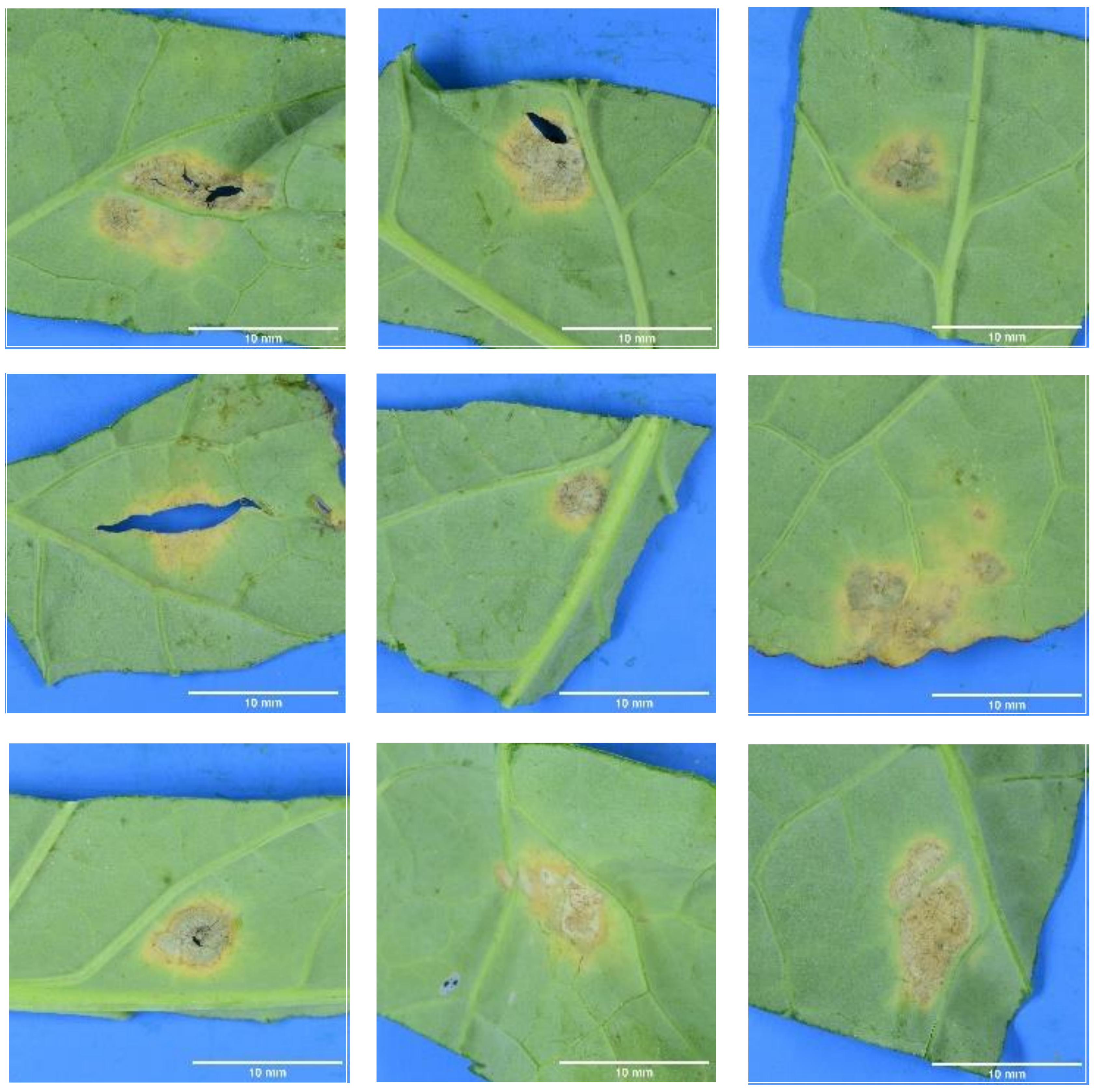
